# Evolution of the genetic architecture of uterine disorders

**DOI:** 10.64898/2026.03.24.713926

**Authors:** Eulalie Liorzou, Hanna Julienne, Maëlle Daunesse, Guillaume Laval, Hugues Aschard, Camille Berthelot

## Abstract

Uterine disorders and menstrual abnormalities are prevalent reproductive conditions with significant clinical consequences. Recent genome-wide association studies have identified hundreds of variants contributing to different uterine disorders, while epidemiological evidence suggests that these disorders co-occur more frequently than expected, implying shared genetic mechanisms. However, the specific genetic variants and biological pathways underlying this shared architecture remain poorly characterized. Here, we conducted a uterus-centric, multi-trait genome-wide association analysis across ten uterine disorders in a European cohort to elucidate this shared genetic architecture. Further, we embedded this architecture in a functional and population genomics framework to investigate its plausible biological mechanisms and its evolutionary history in humans. We confirm strong positive genetic correlations between major uterine disorders and identify 31 independent susceptibility loci jointly affecting the genetic risk of multiple uterine pathologies, substantiating an intertwined biological basis. Populational analyses demonstrated that several of these susceptibility variants exhibit pronounced allele frequency differentiation across global populations suggestive of recent polygenic selection, including variants with well-supported regulatory functions at the *ESR1-CCDC170*, *WNT4*, *SFR1, FOXO1, ITPR1, DMRT1* and *CDKN2B* loci. Notably, we show that most derived alleles acquired during recent human evolution increase risk across multiple uterine disorders and may evolve under antagonistic selection. These findings provide functional annotation and population-level prioritization of genetic variants influencing multiple uterine disorders. They also highlight how past evolutionary histories may contribute to population differences in uterine disease prevalence and pathogenesis.

## Introduction

Menstrual abnormalities are important indicators of reproductive pathological conditions such as endometriosis, endometrial cancers or recurrent miscarriages (Gao et al. 1987; Oehler et Rees 2003; Fraser et al. 2011; Bourdon et al. 2021). Despite their clinical significance, the genetic architecture underlying abnormal menstrual bleeding risk and its association with other uterine disorders remain poorly understood. Over the past 15 years, large-scale genome-wide association studies (GWAS) have identified hundreds of genetic variants associated with different uterine disorders (X. Wang et al. 2022; Buyukcelebi et al. 2024; Rahmioglu et al. 2023; Thibord et al. 2025). For instance, more than 80 loci have been shown to contribute to the polygenic architecture of endometriosis risk (Rahmioglu et al. 2023; Guare et al. 2025; Koller et al. 2025). In parallel, epidemiological studies have reported that several uterine disorders co-occur more frequently than expected (Soliman et al. 2016; Choi et al. 2017; Ye et al. 2022; Fiore et al. 2025; La Vecchia et al. 2025), suggesting that they involve overlapping biological pathways, possibly due to common environmental causes or to shared genetic origins. In support of the latter, the genetic architecture of endometriosis is correlated to that of uterine fibroids (Rafnar et al. 2018a; Gallagher et al. 2019), abnormal menstrual bleeding (Rahmioglu et al. 2023), and endometrial cancers (Painter et al. 2018). Further, this shared genetic architecture across uterine disorders may differ between populations due to their past demographic histories and specific selective pressures. Substantial evidence supports that several uterine disorders show different prevalences among women of African, Asian and European ancestries (Stewart et al. 2017a; Sinharoy et al. 2023; Yamamoto et al. 2017; Kyama et al. 2007; Mecha et al. 2022). For example, uterine fibroids are more frequent and appear earlier in life in women of African ancestry (Stewart et al. 2017b; Giuliani et al. 2020), suggesting population-level differences in genetic susceptibility that may have shaped the risk of uterine diseases across populations. Altogether, genetic and epidemiological reports highlight a shared genetic basis of uterine disorders, although the genetic variants and biological functions involved remain unclear.

In this study, we combine multi-trait GWAS analyses, population genetics and functional genomics to investigate the genetic foundations of shared genetic risk across 10 uterine disorders, and their evolutionary history in the recent human past. Using data from European cohorts, we confirm the presence of strong positive genetic correlations between major uterine disorders, consistent with an intertwined genetic architecture. We identify susceptibility loci showing pronounced allele frequency differences across global populations, including variants with putative regulatory effects at the *ESR1-CCDC170*, *WNT4*, *SFR1, FOXO1, ITPR1, DMRT1* and *CDKN2B* loci. Several of these loci exhibit signatures of recent polygenic selection, suggesting that evolutionary pressures have shaped contemporary patterns of risk and protection in human populations. Altogether, our findings provide a functional and population-level prioritization of genetic variants involved in multiple uterine disorders and illuminate how evolutionary processes may contribute to population differences in uterine disease pathogenesis.

## Results

### Mapping the shared genetic architecture of uterine diseases in European populations

To study the shared genetic architecture of uterine pathologies, we first assessed the estimated genetic correlation captured by GWAS on 10 frequent uterine disorders. We leveraged the FinnGen biobank (Kurki et al. 2023), a Finnish cohort for medical GWAS that has collected phenotypic information for common gynaecological diseases with high numbers of cases and appropriately designed controls for sex specific conditions (i.e. excluding males in this case). We included all traits related to menstrual symptoms and uterine diseases with more than 1,500 documented cases, excluding pregnancy complications (**Methods**), resulting in the inclusion of 10 GWAS summary statistics (**Fig. 1A**, **Table S1**). Using LD score regression (Bulik-Sullivan et al. 2015), we identified significant genetic correlations between 16 pairs of uterine disorders after FDR correction, with correlation coefficients ranking from 0.28 (leiomyoma vs. inflammatory uterine diseases, p-val = 0.024 with BH correction for multiple testing) to 0.94 (excessive frequent and irregular menstruation vs. abnormal uterine bleedings, p-val = 2.10^-49^) (Methods, **Fig. 1B, Table S2**). All disorders except “uterine polyps” and “other menopausal symptoms” shared significant genetic correlations with at least one other trait. These genetic correlations confirm previously reported genetic correlations between endometriosis, leiomyoma and abundant menses in Japanese (Masuda et al. 2020) and European cohorts (Rahmioglu et al. 2023; McGrath et al. 2023), while others are novel (e.g. endometriosis and post-menopausal bleeding), revealing that the genetic burden of common uterine dysfunctions is intricately linked across conditions.

**Figure 1:**
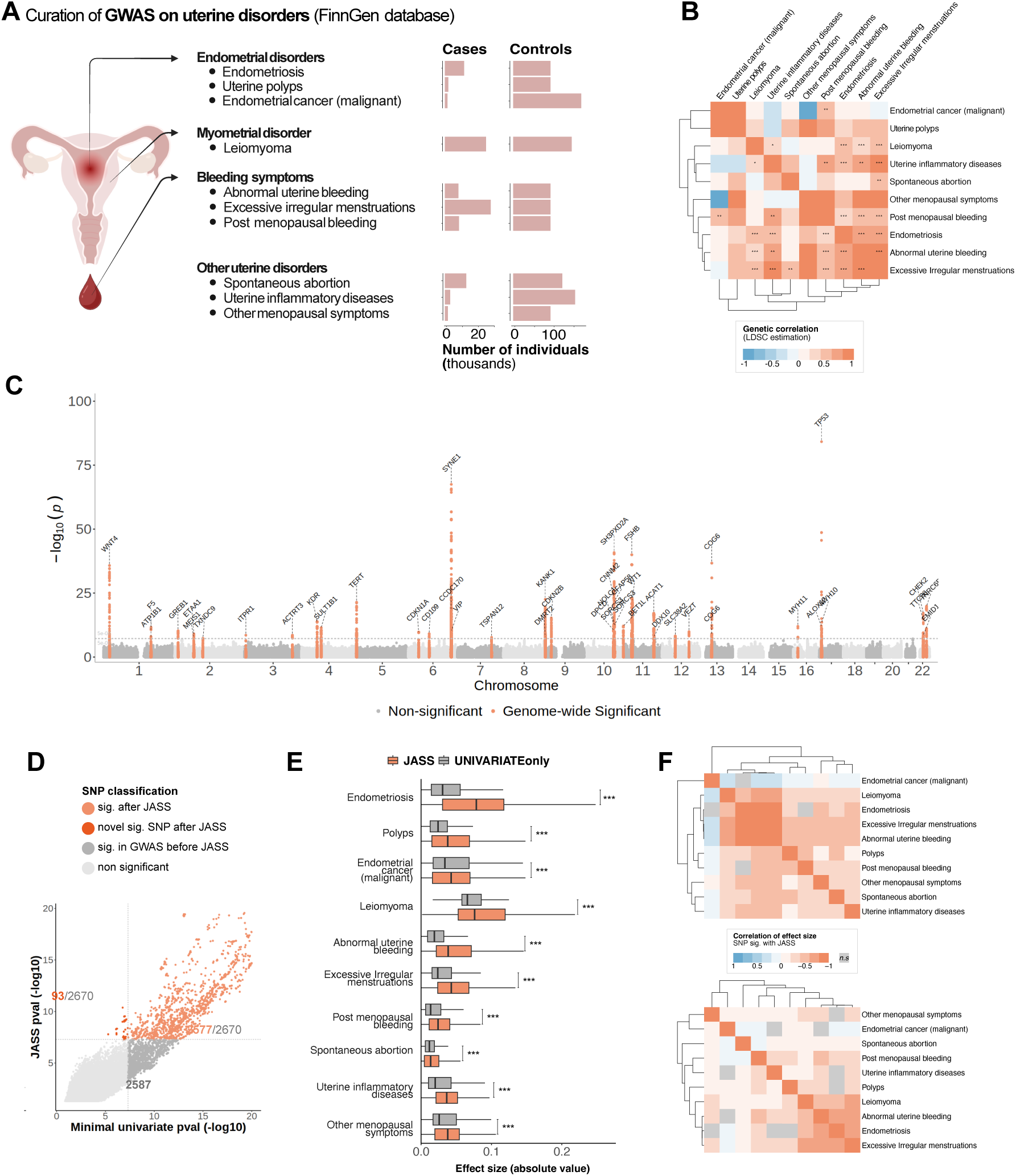
Multi-trait analysis of uterine disorder GWAS. (A) numbers of cases and controls (in thousands) for each of the 10 uterine disorder traits from the FinnGen database included in this study (B). Genetic correlations between traits estimated by LDscore (* p <0.05, ** p <0.01, *** p <0.001). (C). Manhattan plot of the genome-wide significant uSNPs after multi-trait analysis across all 10 uterine disorders (p-val < 5.e-08; significant loci are highlighted in orange). (D). Quadrant plot representing the −log10 p-value of each uSNP after the multi-trait analysis against the lowest p-value across single-trait GWAS. “Sig.” means significant with a p-val < 5.e-08 threshold. The x and y axis are cropped to a maximum −log10 p-value of 20 for readability (uncropped quadrant plot in Fig. S2). (E). Absolute effect sizes (β) of uSNPs significant after multi-trait analysis (uSNPs, orange) and of SNPs significant in single-trait GWAS that do not reach significance in the multi-trait analysis (grey). (F). Correlation of effect sizes for uSNPs (top) and SNPs only significant in single GWAS (bottom) across uterine disorders (Pearson correlation). Colour code represents significant correlations after BH correction for multiple testing; non-significant correlations are shown in grey.

Genetic correlation is a single summary measure that captures the overlap in shared genetic determinants across pairs of traits averaged over the entire genome, and it cannot identify specific genetic loci contributing to the shared genetic risk of uterine diseases. To further identify specific genetic factors shared between common uterine disorders, we analysed those 10 traits using joint analysis of summary statistics (JASS) (Julienne et al. 2020, 2021), a statistical software to conduct multi-trait GWAS association. JASS identified a total of 2,670 variants associated with uterine traits reaching genome-wide significance (p < 5.10^-8^), corresponding to 31 independent genomic loci (**Fig. 1C, Fig. S1, Table S3**). Of these 2,670 pleiotropic SNPs associated with uterine disorders (uSNPs), 93 were not genome-wide significant in any individual uterine disorders GWAS, while 2,577 were significant in at least one single-trait GWAS prior to joint analysis (**Fig. 1D, Fig. S2**). Further, 2,587 SNPs that are significantly associated with at least one individual trait did not reach genome-wide significance in the joint test, suggesting that they contribute less to the shared architecture of uterine disorders. uSNPs prioritized by JASS typically exhibited larger effect sizes (β) across uterine diseases compared to SNPs that were only significant in univariate analyses (**Fig. 1E**). Additionally, uSNPs prioritized with JASS showed overall stronger positive correlations of effect sizes across most uterine diseases compared to SNPs that were only significant in univariate analyses (**Fig. 1F**). Interestingly, jointly prioritized variants showed a strong negative correlation of effect sizes between endometrial cancer and other pathologies, and in particular endometriosis, leiomyoma and menses abnormalities (average r = −0.26; **Fig. 1F**) suggesting potential antagonistic effect of some variants for the risk of endometrial cancer and other uterine diseases.

To replicate this analysis, we used matching uterine traits from the UKBiobank on European ancestry cohorts (Karczewski et al. 2025) (**Methods**, **Table S4**). While the UK Biobank contained fewer cases than FinnGen for most uterine disorders, we found strong directional consistency in the effects of jointly prioritized SNPs for matching traits (**Table S5**). Across uterine diseases, 60% to 83% of uSNPs with a significant association in FinnGen demonstrate a consistent direction of effect in UKBiobank GWAS, which is more than expected by chance (binomial test, p < 0.001 for all comparisons), except for spontaneous abortion and post-menopausal bleeding for which only 43% and 48% of uSNPs have replicated directions of effect, respectively. 583 out of 2,670 uSNPs identified in FinnGen were also significant after joint analysis using UK Biobank uterine disorders GWAS (**Table S6**). These results confirm that we adequately captured SNPs with pleiotropic effects on common uterine diseases in European populations.

### Pleiotropic uterine-associated SNPs are enriched near genes involved in female reproduction, development and senescence

We next investigated how pleiotropic uSNPs associated with uterine disorders may affect gene regulation in the uterus. Almost 98% of uSNPs are non-coding, with 60% (n=1597) falling in introns, 2% in UTRs and 36% (n=962) in intergenic regions (**Fig. 2A, Table S7**), as expected for GWAS significant SNPs. uSNPs are enriched in gene promoters compared to SNPs that are significant in single-trait GWAS only (proportion test; p-val = 4.10^-4^; **Fig. 2B**), and are also enriched in promoter-enhancer contacts captured by Hi-C in endometrial stromal cells (Sakabe et al. 2020) (p-val = 8.10^-6^; **Fig. 2B**). We speculate that uSNPs have more pleiotropic and larger effects on uterine traits in part because they more frequently fall within gene regulatory regions, and in particular gene promoters.

**Figure 2:**
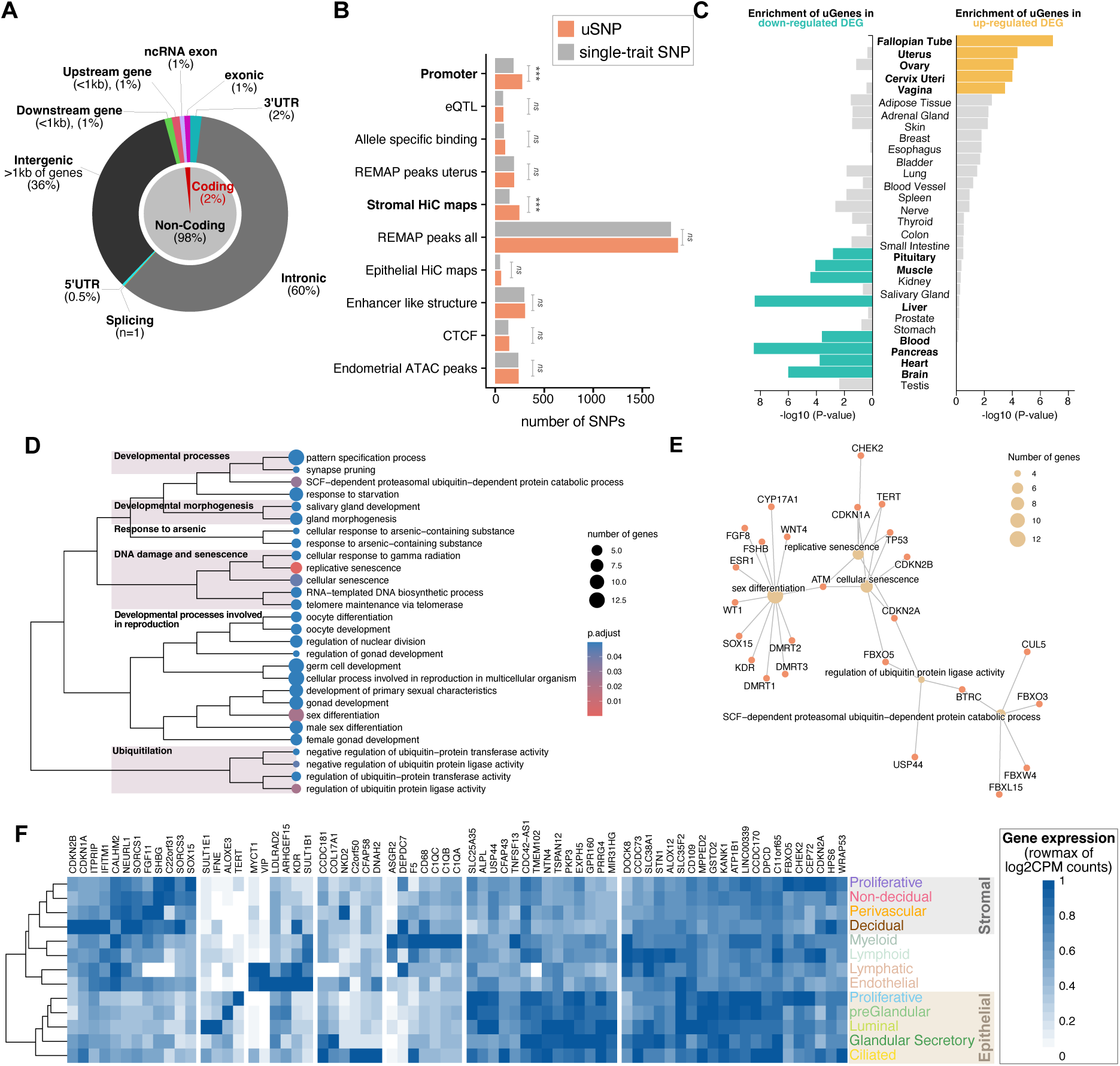
Functional characterisation of uterine-associated SNPs (uSNPs). (A) Distribution of uSNPs across coding and non-coding elements of the genome based on ENCODE annotations. (B) Overlap of uSNPs and single-trait GWAS SNPs (univariate) with functional genomic elements from different databases and from endometrial regulatory profiles (Methods). Absolute numbers of SNPs overlapping the elements are represented (chi-2 test; *n.s* p > 0.05, *** p < 0.001, BH-corrected for multiple testing). (C) Enrichment of uSNPs target genes (uGenes) in organ-specific expression profiles from GTEx. Significant enrichments are represented in colour (turquoise for down-regulated genes, yellow for up-regulated genes, BH-corrected for multiple testing). (D) Gene ontology enrichment analysis using uSNPs target genes. Hierarchical clustering was computed using similarities between GO terms (estimated with Jaccard index). € Network plot of target genes contributing to the 5 first GO enriched terms. Dot size represents the number of target genes associated to each GO term. (F) Gene expression heatmap of uSNPs target genes across endometrial cell types, in log2CPM scaled to the maximum by row. Only genes with cell-type specific expression are represented (full expression heatmap in Fig. S4).

To further explore the functions of genes that may be regulated by uSNPs, we combined functional and location-based information to associate uSNPs with putative target genes (**Methods**, **Fig. S3, Table S7**). In total, uSNPs associate with 227 genes, including well-described loci involved in uterine diseases, endometriosis and uterine fibroids such as *ESR1*, *WNT4*, *WT1* or *FSHB* (Gallagher et al. 2019; Pavlicev et al. 2022; Rahmioglu et al. 2023; Kim et al. 2024; Sheu et al. 2024b). These genes are over-expressed in fallopian tubes, uterus, ovary, cervix and vagina based on GTEX transcriptomes (GTEx Consortium 2017), and under-expressed in several other organs such as brain, blood, muscle or pancreas (**Fig. 2C**; **Methods**). This result highlights that uSNPs are enriched around genes with specialized expression in female reproductive organs.

Gene set enrichment analysis confirms that the putative targets of uSNPs are strongly enriched in terms related to reproductive and developmental processes, involving genes such as *WNT4, WT1, SOX15, NEURL1, FBXO5,* and *CFAP58* (**Fig. 2D-E**). The strongest enrichment was found for genes associated with replicative senescence (38-fold enrichment; **Fig. 2D, Table S8**), driven by canonical regulators such as *TERT, TP53, CDKN1A, ATM* and *CHEK2* (**Fig. 2E**). Additionally, we find enrichment in terms related to ubiquitylation (e.g. *BTRC, CDKN2A, FBXO5*) and arsenic response (e.g. *GSTO1, GSTO2, SLC38A2*). While those three processes are not specific to uterine or reproductive physiology, they represent conserved cellular networks that may indirectly influence reproductive function. This suggests that pleiotropic uterine SNPs influence both canonical reproductive pathways and broader cellular processes, highlighting their potential role in linking systemic biological networks with female reproductive functions.

To investigate how uSNPs might influence uterine cell functions, we re-analysed endometrial single-nuclei transcriptomes of healthy donors from the HECA atlas (Marečková et al. 2023), (**Fig. S4A**). This showed that 74% of uSNP target genes are expressed in at least one uterine cell type (**Fig. S4B**). Many of these genes, including well-documented loci such as *ESR1, WT1,* and *WNT4*, are broadly expressed across multiple uterine cell types and likely have ubiquitous effects across the reproductive system. More interestingly, several target genes displayed cell-type–specific expression (**Fig. 2F**). For example, *CDKN2B, CDKN1A, ITPRIP* and *IFITM1* were preferentially expressed in decidual stromal cells, while *CCDC181, COL17A1, CFAP58* and *DNAH2* were strongly expressed in ciliated epithelial cells. This observation suggests that uSNPs may also capture pleiotropic effects resulting from altered functions in restricted cell populations.

As expected for pleiotropic loci, most uSNP target genes were also expressed in tissues beyond the uterus. Among the few targets not detected in healthy uterus, several showed highly tissue-specific expression in the testis, pituitary or brain (e.g., *FSH, INA, CYP17A1*) (**Fig. S5**). These patterns suggest that some uSNPs may exert pleiotropic effects on uterine traits via systemic endocrine or neuroendocrine pathways, or that certain genes may become aberrantly expressed in the uterus under pathological conditions.

### Substructure of uSNPs with risk-increasing or protective effects on uterine disorders

We next investigated how the shared genetic architecture of uterine diseases is structured into groups of variants with similar effects on multiple disorders. To obtain a set of largely independent variants, we reduced the full set of 2,670 uSNPs to 235 lead uSNPs in pseudo-independent linkage disequilibrium blocks (R^2^ < 0.5), and identified the ancestral and derived alleles for each lead uSNP (**Methods**). Then, we performed unsupervised K-means clustering on effect sizes (β) of lead uSNP derived alleles across traits (**Methods**). This analysis identified an optimal clustering with two main clusters representing uSNPs for which the derived allele has an overall protective effect (negative effect size) across most uterine disorders (cluster 1), or has an overall risk-increasing effect (cluster 2; positive effect size) (**Fig. 3A, Table S9**). Risk-increasing cluster 2 contained twice as many lead uSNPs as protective cluster 1, suggesting that recently-acquired alleles are more frequently deleterious in terms of uterine disorder risk.

**Figure 3:**
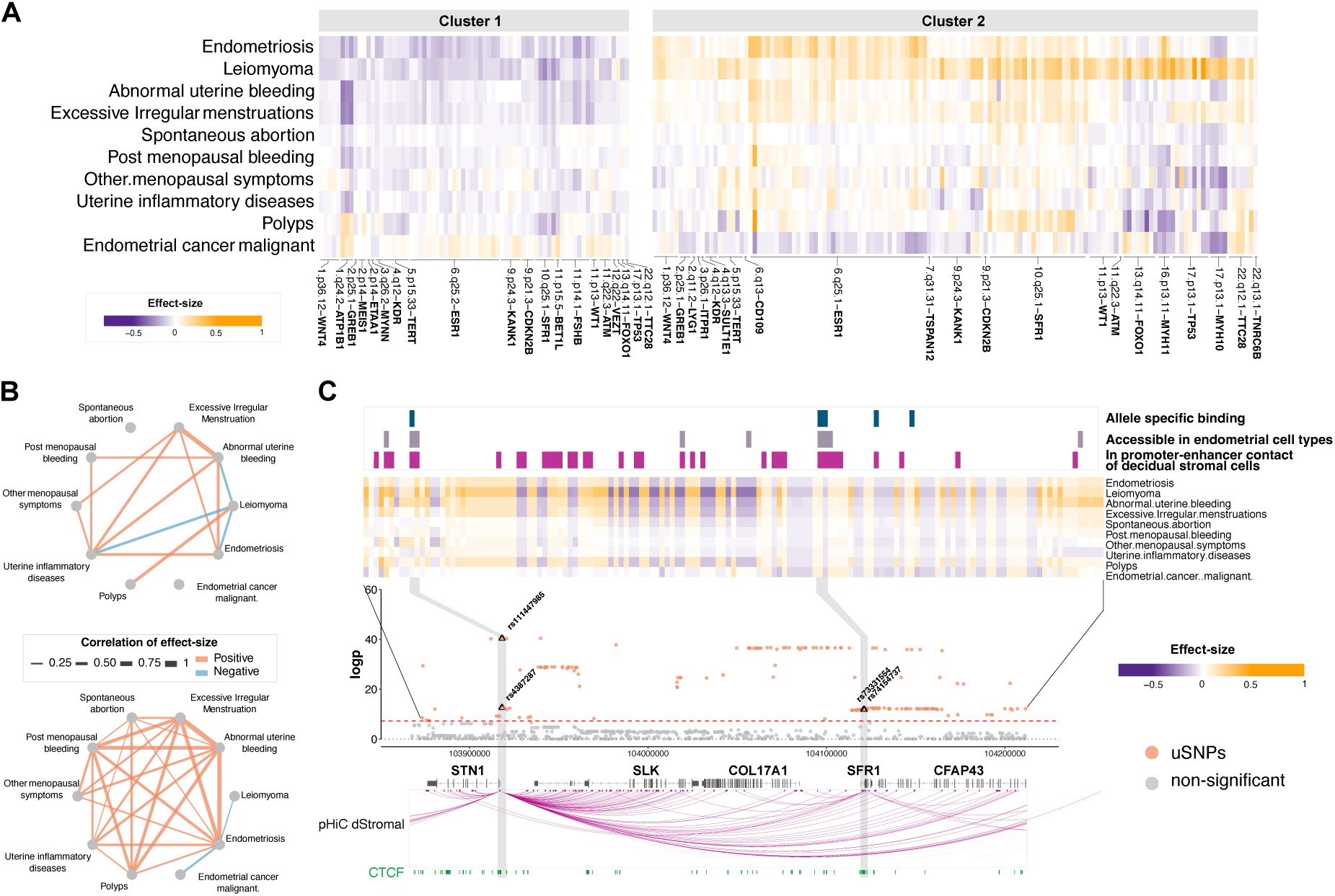
Substructure of uSNPs effect sizes across uterine disorders. (A) Heatmap of effect sizes for derived alleles of pseudo-independent uSNPs across uterine traits. Cluster 1 and cluster 2 were determined by k-means clustering on uSNPs effect sizes. For each cluster, uSNPs are ordered by genomic position and labelled by their nearest gene name. (B) Correlations of uSNPs effect sizes across uterine traits for cluster 1 (top) and cluster 2 (bottom). Only significant correlations are represented (BH-corrected for multiple testing). (C) Representation of the *SFR1* locus showing normalised effect sizes of derived alleles and functional annotations of uSNPs.

To further explore how protective and risk-increasing loci associate with different disorders, we computed pairwise correlations of normalized effect sizes between disorders for each cluster (**Fig. 3B**). Variants in cluster 1 had correlated and protective effects on endometriosis and different menses and menopause disorders, but negative correlations and deleterious effects with other traits such as endometriosis and leiomyoma – suggesting that these variants tend to be protective for either disease but not both. Variants in cluster 2 had overall correlated risk-increasing effects on most disorders, with the notable exception of malignant endometrial cancer and leiomyoma, which were negatively correlated with endometriosis. Specifically, we identified an antagonistic pleiotropic relationship between endometriosis and malignant endometrial cancer, where derived alleles at lead uSNPs in cluster 2 typically increased risk for endometriosis but had overall protective effects on endometrial cancer, in particular at the *ESR1* and *GREB1* loci (**Fig. 3A-B**). By focusing on a subset of largely independent SNPs that associate with multiple disorders, this analysis breaks down the genome-wide genetic correlations identified in **Fig. 1B** into more nuanced components with correlated or antagonistic pleiotropic effects between disorders.

This analysis highlighted a locus of particular interest on chromosome 10 (10.q25.1), containing a remarkably high concentration of pleiotropic uSNPs of large effect sizes on multiple disorders, especially leiomyoma, excessive irregular menstruation and polyps (**Fig. 3C**). This locus is rich in putative non-coding regulatory elements active in the endometrium (accessible in single-cell ATAC-seq data from human endometrium (Vrljicak et al. 2023), and corresponding to promoter-enhancer contacts in decidual stromal cells assessed by promoter-capture Hi-C (Sakabe et al. 2020)). rs4387287*C and rs111447985*\**C, which overlap with CTCF binding sites and are predicted to enhance the binding of the BRD4 transcription factor, respectively, are notable in this locus. Both SNPs are situated within the promoter region of the STN1 gene, a chromatin region that is accessible across various endometrial cell types (**Fig. 3C**, **Table S7**). Remarkably, these SNPs coincide with 116 Hi-C contacts in decidualized endometrial stromal cells, establishing connections notably with the promoters of *SFR1*, *CFAP43*, and *ITPRIP*, suggesting robust regulatory activity of this chromatin region across endometrial cell types. In healthy endometrium, the *STN1* gene demonstrates consistent expression across different endometrial cell types, with particularly elevated levels in endometrial glandular cells and immune cells (**Fig. 2E**). *STN1* is known to play a crucial role in cancer progression, acting as an upstream regulator of the epithelial-to-mesenchymal transition (Nguyen et al. 2023; Dong et al. 2025). Notably, previous GWAS studies have associated this locus with the risk of endometriosis (Sheu et al. 2024a), and our findings suggest that variants in this locus may have pleiotropic effects on other uterine disorders, such as uterine leiomyomas. While the functional roles of these candidate uSNPs in endometrial cells await experimental validation, they have the potential to disrupt the regulatory landscape of *STN1* expression, as well as those of *SFR1*, *CFAP43*, and *ITPRIP*.

### Evidence of polygenic selection at uterine disease risk loci in European populations

Because genetic variants affecting reproductive functions may be particularly accessible to natural selection due to their direct effect on fitness, we next investigated whether the shared genetic architecture of uterine disease risk displays evidence of polygenic selection in human populations(Pritchard et al. 2010; Pritchard et Di Rienzo 2010; Stephan et John 2020). We hypothesized that uSNPs may be under stronger evolutionary pressures than background SNPs because they may have pleiotropic effects on gene expression and therefore be more likely to evolve under selection (Pavlicev et Wagner 2012; Hämälä et al. 2020). We compared the fixation index (F_st_) distributions between the 191 lead uSNPs and 1,000 sets of control SNPs matched for recombination rate, minor allele frequency (MAF) and local gene density, in populations from the 1000 Genomes Project (The 1000 Genomes Project Consortium et al. 2015) (**Methods**). F_st_ measures allelic differentiation between pairs of populations and is increased for variants that experienced past positive selection (Holsinger et Weir 2009; Stephan 2016). The F_st_ distribution of lead uSNPs is shifted towards higher F_st_ values in the reference European population (CEU) compared to the reference East Asian (CHB) and African (YRI) populations (**Fig. 4A**; CEU vs. CHB, p-val = 0.024, CEU vs. YRI, p-val = 0.0009; Wilcoxon test with BH correction for multiple testing). In contrast, we detected no significant difference in F_st_ values between YRI and CHB populations. We next compared the integrated haplotype scores (iHS) of lead uSNPs and controls in each population, which measures the local strength of selection based on haplotype homozygosity to detect more recent selection events (Voight et al. 2006a). uSNPs have overall higher absolute iHS than matched controls in Europeans (CEU), although the signal does not reach significance after multiple testing correction (**Fig. 4B**; p-val = 0.087; Wilcoxon test with BH correction for multiple testing). These results suggest that uterine disease risk loci captured from European cohorts have likely been influenced by past selective pressures resulting in polygenic allelic differentiation between human populations.

**Figure 4:**
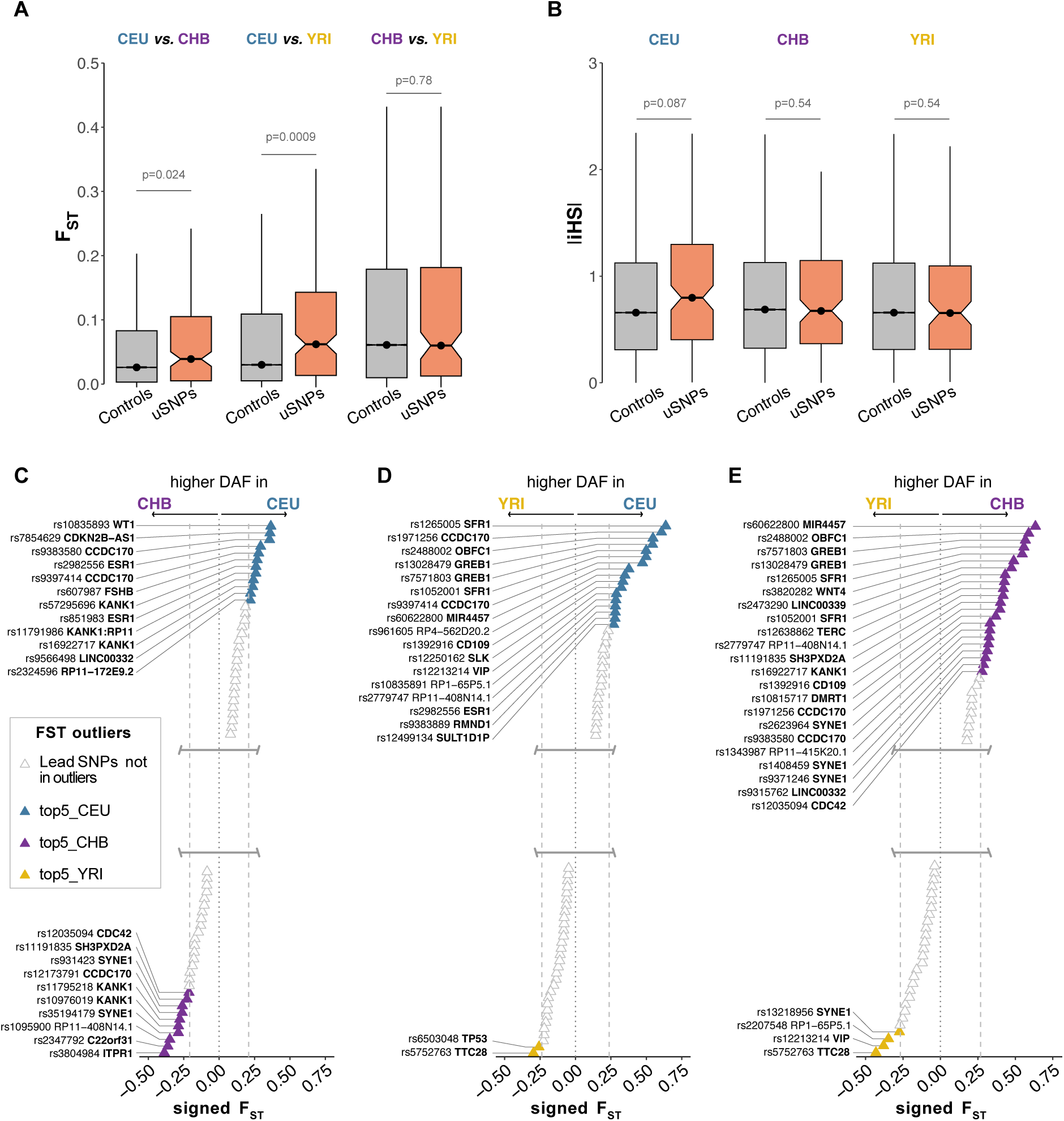
Evolutionary statistics of uSNPs in reference European, African and East-Asian populations. (A-B) Distributions of F_st_ (A) and iHS values (B) at pseudo-independent lead uSNPs compared to 1,000 sets of matched control SNPs (Wilcoxon rank sum test, BH-corrected for multiple testing). (C-D-E) Signed F_st_ values of the top pseudo-independent lead uSNPs for all three population comparisons: (C) CEU (Northern European from Utah) vs. CHB (Chinese Han from Beijing); (D) CEU vs. YRI (Nigerian Yoruba from Ibadan); (E) YRI vs. CHB. uSNPs in the top 5% genome-wide F_st_ values are represented in colour; signed F_st_ scores point towards the population where the derived allele frequency is the highest.

To highlight specific uSNPs of interest that might have experienced stronger events of selection, we next focused on uSNPs with outlier F_st_ values (top 5%), which exhibit very strong allelic differentiation between populations and are prime functional candidate loci, similarly to previous approaches (Quach et al. 2016). We find that 50 uSNPs out of 191 uSNPs tested are F_st_ outliers in at least one comparison (23%, p-val = 0.0004, binomial test, **Table S10**, **Methods**). Moreover, uSNPs are over-represented in F_st_ outliers in both the CEU vs. CHB and CHB vs. YRI comparisons (**Fig. 4C-E**; CEU vs. CHB, p-val < 2.2.10^-16^; CEU vs. YRI, p-val = 0.12; CHB vs. YRI, p-val = 0.01; empirical test with BH correction for multiple testing, **Methods**). We compared the GWAS effect sizes of uSNPs with outlier F_st_ values to non-outlier uSNPs and found no significant differences between groups (**Fig. S7**).

Amongst uSNPs with high F_st_ values, we prioritized a list of candidate variants with putative functional impact on uterine cell types as described in **Fig. 2**. This analysis highlights 17 uSNPs with both evidence of allelic differentiation across human populations and functional importance for uterine cell functions, spanning the *ESR1*, *WNT4*, *SFR1*, *FOXO1*, *KANK1* and *CDKN2B* loci (**Table 1**). As *CDKN2B*, *ITPR1* and *WNT4* are mostly expressed in endometrial stromal cells, and *DMRT1* and *CCDC170-ESR1* in epithelial cells, we speculate that variants at these loci may mostly affect these cell types. Interestingly, variants at the *SFR1* locus, and presented in **Fig. 3C**, also show evidence of allelic differentiation across populations and display the highest F_st_ score of all tested uSNPs (**Fig. 4D**). Additionally, rs3820282 at the *WNT4* locus was recently highlighted as a pleiotropic variant with beneficial roles in gestation length and pre-term birth while increasing the risk of endometriosis, breast cancer and ovarian cancer (Pavlicev et al. 2022). Functional validation in human cell lines and a transgenic mouse model confirmed that rs3820282 increases the binding of ESR1 and induces an over-expression of *WNT4* in endometrial stromal cells (Pavlicev et al. 2022).

**Table 1:**
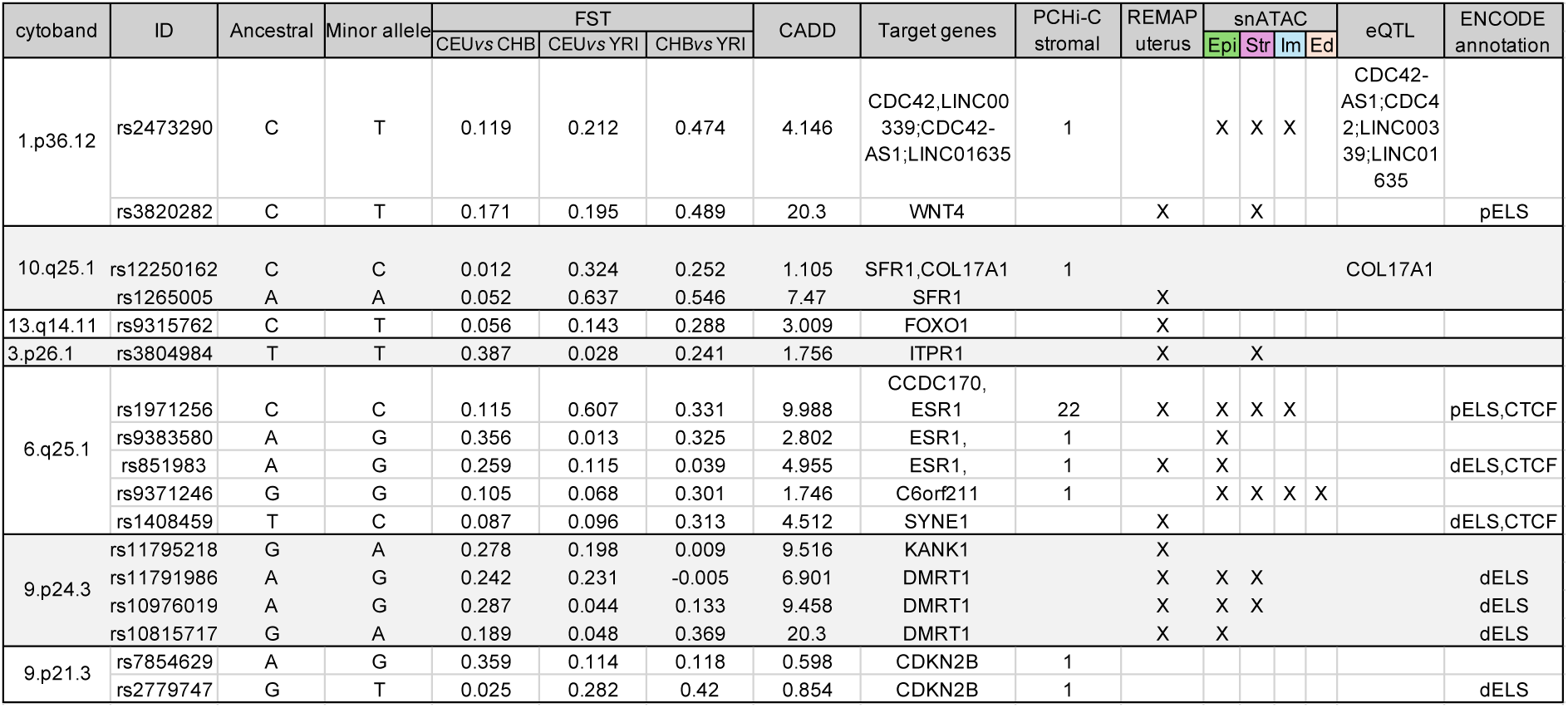
Prioritized susceptibility loci harbouring candidate uSNPs with signatures of strong allelic differentiation and with regulatory potential in endometrial cells. The full table of functional and evolutionary annotation of those variants is available in **Table S10**. pELS: proximal enhancer-like structure; dELS: distant enhancer-like structure; Epi: epithelial; Str: stromal; Im: immune; Ed: endothelial.

Altogether, our results support that a sizeable fraction of uSNPs display evolutionary signatures of positive selection in European populations, and further prioritize candidate SNPs within these genomic regions with joint evolutionary and functional signatures.

### The *ESR1-CCDC170* locus displays strong evidence of positive selection in European populations

Among genetic loci with evidence of recent selective pressures in human populations, the genomic region including both the *CCDC170* and *ESR1* genes is of particular interest as it carries a high number uterine-relevant SNPs with outlier F_st_ values (**Table 1**, **Fig. 5A**). Notably, rs1971256 displays the strongest F_st_ value of this locus (CEU vs. YRI, F_st_ =0.607; CHB vs. YRI, F_st_ = 0.331) and the highest score of pathogenicity (CADD = 9.988, **Table 1, Methods**). Additionally, this locus exhibits well-supported evidence of selective sweeps in the last 100,000 years across all European populations and in one African population, which constitute rare events of strong positive selection (Laval et al. 2021) (**Fig. 5C**). Indeed, we observe that the rs1971256 derived allele (T) has become the major allele in most human populations, except those of African descent (**Fig. 5D**). This variant was previously prioritised as increasing the risk of endometriosis as well as endometrioid and clear cell ovarian cancers (Sapkota et al. 2017a; Mortlock et al. 2022), but its functional impact on endometrial cell types has not been investigated. In this study, we found that the derived allele of rs1971256 has a pleiotropic protective role across multiple uterine traits, notably endometriosis, abnormal menstrual bleeding, excessive and frequent menstruation and inflammatory uterine diseases (**Fig. 5B)**. This variant is in the promoter of *CCDC170*, and additionally displays 22 significant Hi-C contacts with the *ESR1* promoter and introns identified in decidual stromal cells (**Fig. 5C**). Additionally, this region is in accessible chromatin across several endometrial cell types and is a predicted CTCF binding site by ENCODE (Dunham et al. 2012) (**Fig. 5C**). In healthy endometrium, *CCDC170* is particularly expressed in proliferating endometrial epithelial cells, pre-glandular cells and ciliated cells, and *ESR1* is expressed across most cell types during the proliferative phase (**Fig. 5E**). Other uSNPs with outlier F_st_ value in this locus are accessible in endometrial epithelial cells preferentially (**Table 1**), which additionally suggests that this genomic locus might contribute to uterine disease pathogenesis by affecting endometrial epithelial cells. Future functional studies may reveal how these genetic variants that have reached higher frequencies across many human populations affect the physiology of endometrial cells and protect against endometriosis or abnormal uterine bleeding.

**Figure 5:**
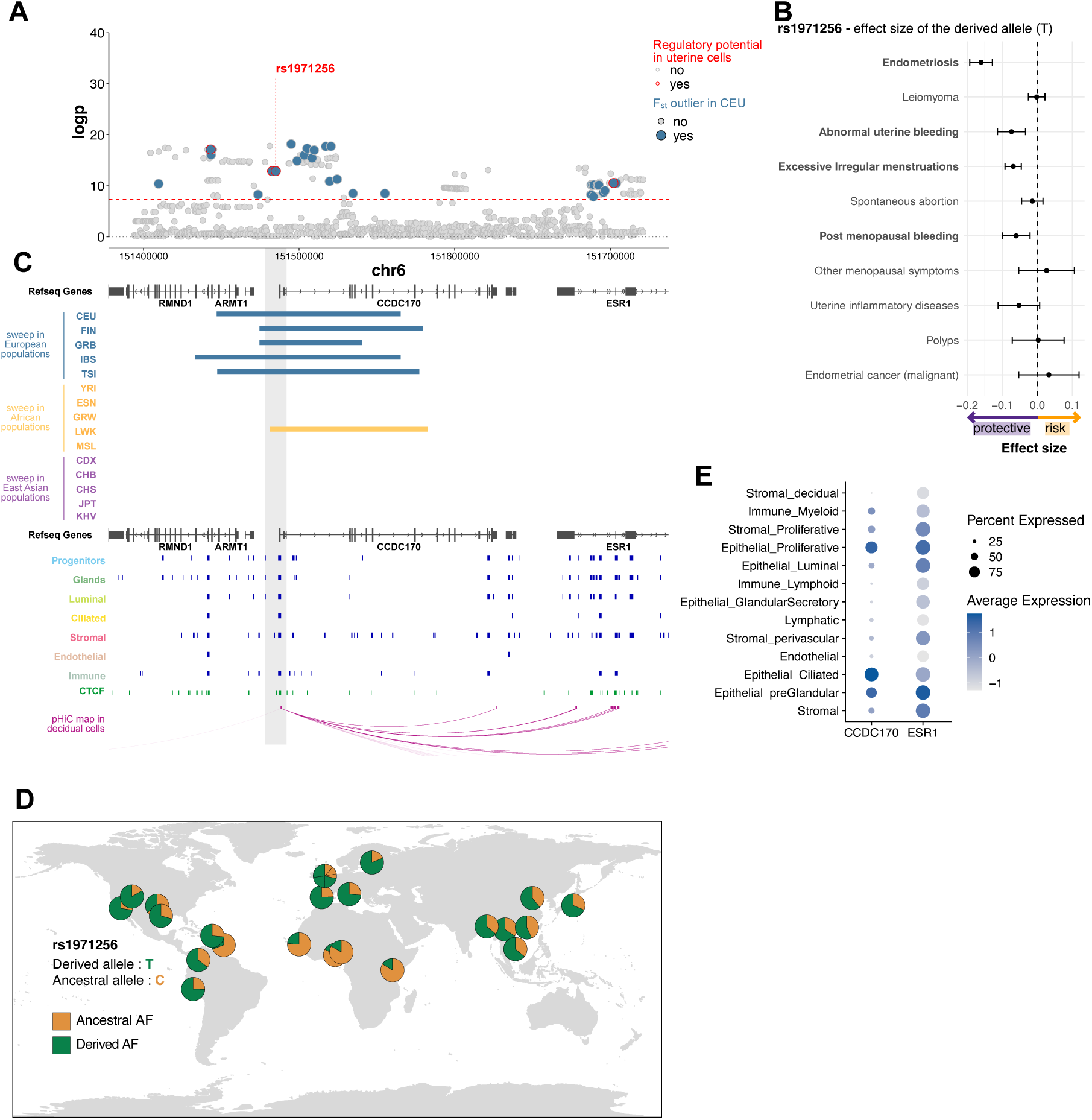
Evolutionary and functional annotation of the *CCDC170-ESR1* locus. A. Regional plot highlighting significant uSNPs with outlier F_st_ values in the CEU population (in blue) and with functional evidence of regulatory activity in endometrial cells (circled in red). B. Heatmap of the effect sizes of the derived allele at rs1971256 across uterine disorders. C. Annotation of selective sweeps at the *CCDC170-ESR1* locus from Laval et al. 2021 across 1000 Genomes populations, overlaid with open chromatin peaks identified across endometrial cell types, CTCF binding sites from ENCODE and promoter Hi-C contacts captured in decidual stromal cells. D. Global map of rs1971256 allele frequencies across populations included in the 1000 Genomes Project. E. Average gene expression of *CCDC170* and *ESR1* across endometrial cell types using single-nuclei RNA-seq from healthy human endometrium.

## Discussion

In this study, we explored how combining uterus-centric GWAS summary statistics can illuminate shared genetic and cellular mechanisms of uterine dysfunction. We integrated the polygenic architecture of ten uterine disorders – most of them associated with menstrual symptoms – and confirmed extensive, genome-wide genetic correlations across these disorders. Further, we prioritized 31 major susceptibility loci with pleiotropic effects contributing to this shared genetic burden and potentially involved in the pathogenesis of multiple uterine disorders based on functional and evolutionary evidence. Compared to previous work investigating genetic correlations between pairs of diseases (Painter et al. 2018; Rafnar et al. 2018a; Gallagher et al. 2019; Masuda et al. 2020; Rahmioglu et al. 2023), this approach integrates evidence across multiple uterine pathologies into a unified statistical framework to reveal genetic loci impacting core pathways within the uterus. To date, few studies have investigated how organ-centric integration of GWAS statistics can highlight genetic variants of functional significance to organ function and dysfunction (Sheng et al. 2021; Levin et al. 2022; J. Song et al. 2024; X.-Y. Wang et al. 2025).

The substantial genetic correlations observed amongst uterine disorders are consistent with prior evidence that gynaecological disease susceptibility frequently involves overlapping biological pathways broadly related to senescence, hormonal response and reproductive development mechanisms (Sapkota et al. 2017b; Rafnar et al. 2018b; Bulun et al. 2019; Cardoso et al. 2020; Bulun et al. 2025). Further, our results show that most genetic variants identified by our approach have congruent effects and either decrease or increase risk across diseases, consistent a fundamental aetiology common to uterine disorders. One exception was endometrial cancer, which was typically anticorrelated with other uterine disorders, capturing antagonistic genetic effects. A major hurdle to understand disease mechanisms underlying this shared genetic risk is that most GWAS variants are likely non-functional or their functional potential is rarely investigated in detail. Here, to support biological interpretation, we integrated a broad array of functional genomics resources to clarify how these variants may impact gene expression across tissues and specifically in the uterine endometrium. Our results reveal that pleiotropic variants affect loci harbouring genes whose expression is highly biased towards female reproductive tissues. Further, most of these genes are highly expressed in endometrial cells, confirming that this multi-trait analysis strategy captures variants within loci of direct interest for uterine biology. Our results suggest that at least a subset of shared susceptibility loci act by disrupting gene regulation in different uterine cell types. Although functional annotation alone cannot establish causality, the convergence of genetic associations, regulatory context in uterine cells, and uterine expression of nearby genes strengthens the plausibility that these loci contribute mechanistically to uterine disease pathogenesis, rather than reflecting indirect systemic effects. However, experimental validation will be required to resolve causal variants and target genes and confirm the functional consequences of these variants.

To understand how mutations that arose recently in human history contribute to uterine disorder risk, we investigated the directionality and effect sizes of derived alleles at pleiotropic loci. To our knowledge, this is a novel approach as most studies define reference and alternative alleles arbitrarily and therefore rely on absolute effect sizes only, ignoring directionality of effects. Considering the risk contributions of ancestral and derived alleles revealed that recently acquired alleles were almost twice more likely to increase disease risk. Overall, our findings support that most recent mutations in the human genome that impact uterine functions are deleterious. The relatively high frequencies at which some of these derived alleles are present in human populations suggests that either this deleterious effect is too weak to be purged by natural selection, or these variants may have been influenced by antagonistic selection acting on other genetically correlated traits.

To further investigate how these pleiotropic variants have been influenced by past selective pressures, we considered different metrics of natural selection based on allelic frequencies and haplotype homozygosity in populational data (Weir et Cockerham 1984; Voight et al. 2006b; Pritchard et al. 2010). Our results support that pleiotropic variants acting on uterine disorder risk present signatures of selection in European populations. This signal was stronger when using the F_st_ score compared to the iHS score, which detects independent events of rapid selection (Voight et al. 2006b), suggesting that allelic differentiation at these loci more likely reflects polygenic selection. We however note that alternative explanations such as demographic history, background selection, or genetic drift cannot be entirely excluded. Moreover, the modest number of loci identified in our study (31 loci represented by 235 pseudo-independent variants) limits statistical power relative to large-scale studies of polygenic adaptation in traits such as height or metabolic phenotypes (Berg et al. 2019; Choin et al. 2021; Kun et al. 2023). Nonetheless, evidence of consistent allelic differentiation across multiple loci suggests that uterine disorder genetic risk has been shaped by recent population history, and that the risk architecture captured in European populations may not reflect that of other populations, highlighting the importance of expanding GWAS studies to population samples of diverse genetic ancestries.

Our observations overall suggest that, despite being on average deleterious, pleiotropic variants impacting uterine disease risk tend to be selected by polygenic adaptation processes. This ties in with theories of antagonistic selection at pleiotropic loci, where pleiotropic variants are adaptative for early-life traits and reproductive success but become deleterious in later life stages, particularly by increasing disease burden (Rose 1982; Pavlicev et Wagner 2012). This interpretation is supported by observations that some uterine disorders are genetically correlated with traits related to fertility, such as age at menarche, or to pregnancy (Rahmioglu et al. 2023; Benonisdottir et al. 2024; Venkatesh et al. 2025; Pujol Gualdo et al. 2025). Previous work has reported that variants affecting reproductive traits are often located near genes involved in cellular aging and lifespan regulation (Rodríguez et al. 2017a; D. Wu et al. 2022), which we also confirm here with an enrichment of senescence-related genes close to the genetic variants prioritized by our approach. Human genetic studies increasingly document genetic correlations between reproductive traits and later-life disease risk, although the molecular mechanisms underlying these relationships remain poorly characterized (D. Wu et al. 2022; Chen et Zhang 2020; Rodríguez et al. 2017b).

Our study comes with a number of limitations. First, we did not evaluate associations between prioritized variants and other reproductive phenotypes such as age at menarche, age at first birth or gestational duration, as those traits are not included in the Finngen biobank. However, several of these traits have previously shown evidence of selection favouring earlier reproductive maturation (Lahdenperä et al. 2004; W. Song et al. 2021; Long et Zhang 2023; Xiang et al. 2024). Future work integrating multi-trait GWAS across reproductive life-history traits may clarify whether recently acquired alleles increasing uterine disorder risk are also associated with reproductive advantages. Another limitation of this work is that GWAS data used in this study were derived from individuals of European ancestry, limiting the generalizability of both pleiotropic associations and population genetic analyses. Additionally, diagnostic heterogeneity across uterine disorders in biobank-based phenotyping likely includes cases misclassified as controls, which likely reduces power to detect significant effects for some traits. Indeed, while the Finngen biobank includes more uterine disorder cases compared to other biobanks (Kurki et al. 2022; Constantinescu et al. 2022), the proportions of cases vs. controls remain lower to the estimated prevalence for several uterine disorders in European populations. For instance, uterine fibroids have an estimated prevalence of ∼70% in women after 50 years (Stewart et al. 2017b), but only represent 17% of patients in the Finngen cohort.

In summary, our study explores the shared genetic architecture of uterine disorders and identifies 31susceptibility loci with pleiotropic effects, several of which show marked population differentiation and regulatory activity in uterine tissues. These findings pave to the way to explore common biological pathways underlying multiple gynaecological conditions and suggest that some components of uterine disease risk are shaped by population-specific genetic histories. Expanding multi-trait and evolutionary analyses to larger number of traits related to female reproductive functions and to cohorts from diverse ancestry, will be critical for clarifying the generality and biological significance of these genetic variants.

## Material and Methods

### Selection of GWAS on uterine disorders in the Finngen database

To explore the genetic architecture of uterine disorders, we leveraged available summary statistics from FinnGen, a hospital cohort with well-defined phenotypes uterine disorders, including sex-specific designs for male/female-specific traits, and which includes higher case counts for uterine disorders compared to other publicly accessible cohorts such as the UKBiobank (See **Table S1** and **Table S4**). FinnGen database (r.7) was used to select GWAS summary statistics related to uterine traits and diseases using “uterine”, “endometrium”, “menstruation”, “fertility”, “menarche”, “menopause”, “leimyoma” and “bleeding” as keywords. We did not include traits and diseases related to pregnancy complications as the focus of our study concerns uterine functions outside of pregnancy. Phenotypes with fewer than 1,500 cases were filtered out. Endometriosis included a number of subcategories, and we only retained the broad phenotype with the largest number of cases (“endometriosis”), resulting in 10 final uterine phenotypes: endometriosis (ICD10-N80), malignant endometrial cancer (ICD10-C54), leiomyoma (ICD10-D25),, abnormal uterine bleeding (ICD10-N93), inflammatory uterine disease (ICD10-N71), post-menopausal bleeding (ICD10-N95.0), other menopausal disorder (ICD10-N95.8), excessive-frequent-irregular menstruation (ICD10-N92),, uterine polyps (ICD10-N84.0), and spontaneous abortion (ICD10-O03) (**Table S1**). We retained “spontaneous abortion” as a proxy for uterine receptivity, as failure in effective endometrial preparation to implantation largely contributes to miscarriage in humans (Lucas et al. 2016; Muter et al. 2021), although embryo defects also contribute. Additionally, we also retained post-menopausal bleeding and other menopausal disorders, which can be symptoms of post-menopausal uterine diseases, notably endometrial cancer or uterine leiomyomas (Makker et al. 2021; Munro 2019).

### Selection of GWAS on uterine phenotypes in Pan-UK Biobank database

For replication purposes, we selected uterine-associated phenotypes in the Pan-UK Biobank matching those selected from Finngen and using the same inclusion criteria (*See* **Table S4**). Summary statistics performed on individuals of European ancestry were downloaded from the Pan-UK Biobank website, resulting in the inclusion of 8 uterine phenotypes: malignant endometrial neoplasm (ICD10-C54), leiomyomas (ICD10-D25), endometriosis (ICD10-N80), uterine polyps (ICD10-N84.0), irregular menstrual cycle bleedings, excessive menstrual cycle bleeding, post-menopausal bleeding, and miscarriage. Inflammatory uterine diseases and other menopausal disorders of the uterus were not phenotyped in the Pan-UK Biobank.

### Summary statistics curation and pre-processing

GWAS summary statistics were harmonized using the JASS pre-processing pipeline (Julienne et al. 2020). This pipeline consists in (i) harmonizing heterogeneous formats of summary statistics, (ii) aligning the effect allele of each study to the effect allele of the reference panel, (iii) removing strand-ambiguous variants and variants with low sample sizes, (iv) computing trait heritability, and genetic and residual covariances across traits using the LDscore regression. For FinnGen summary statistics, we used a Finnish reference panel and LDscore data derived from GnomAD v2 (https://gnomad.broadinstitute.org/downloads#v2-linkage-disequilibrium). For Pan-UK Biobank summary statistics, we used the European reference panel and LDscore data based on the 1000G database provided as part of the JASS pipeline. We used positional SNP identifiers (defined as the combination of chr:pos:ref:alt) to map all summary statistic SNPs to the reference panel. For both datasets, no imputation step was performed. The summary information for each summary statistics used as input for the JASS preprocessing pipeline can be found in **Table S1** and **Table S4**. We report the genetic correlation and the significance of the genetic correlation computed by LDscore (B. K. Bulik-Sullivan et al. 2015; B. Bulik-Sullivan et al. 2015) and corrected for false discovery rate using the Benjamini-Hochberg correction (**Table S2**).

### Joint analysis of uterine trait summary statistics with JASS

Multi-trait analysis of uterine traits summary statistics was conducted using the omnibus test implemented in the JASS suite (Julienne et al. 2020). JASS output summary statistics were further processed by associating each SNP to their RSID using the initial summary statistics from FinnGen. For the 18,524 SNPs without an RSID in initial summary statistics, we used ANNOVAR (hg38) SNPs RSID and retrieved RSIDs for an additional 4,680 SNPs. The remaining 14,044 SNPs without official RSIDs in Annovar were named using their positional identifier.

For each SNP, we reported the p-value of the joint test and the smallest p-value across univariate GWASs. The ability of JASS to capture variants with pleiotropic effects across uterine traits was investigated (1) for each uterine trait independently, by comparing the effect sizes of uSNPs vs single-trait SNPs; and (2) across uterine traits, by computing pairwise uSNPs effect size correlations. We also compared effect size magnitudes of uSNPs and single-trait SNPs for each uterine trait independently using Wilcoxon rank sum tests, FDR-corrected using the Benjamini-Hochberg correction.

### Replication analysis using GWAS from the Pan-UK Biobank

For each trait, we compared the directionality of allele effects between effects estimated in the Finngen cohorts and in the Pan-UK Biobank cohorts. For all uSNPs identified with the Finngen dataset and also present in Pan-UK Biobank summary statistics, we tested whether directions of effects were consistent after matching effect alleles between both biobanks. Using a directional binomial test, we tested whether the proportion of uSNPs with replicated effect directions was greater than expected by chance (**Table S5**). After grouping uSNPs into loci, we additionally tested if the number of loci with replicated effect directions was greater than expected by chance using the same approach (**Table S5**).

### Selection of independent and lead SNPs with FUMA

FUMA (v1.5.2) was used to define 241 lead SNPs with a linkage disequilibrium correlation r^2^ < 0.5, which is the default threshold used by PLINK (Purcell et al. 2007). Lead SNPs within 500 kb of each other were merged as a single genomic locus, resulting in the identification of 31 genomic risk loci. To calculate r^2^, MAF (minor allele frequency) and conduct LD analyses, European population genetic data from the 1000G Project phase 3 (2015) was used as the reference panel.

### Functional annotation of candidate SNPs

The ANNOVAR (v2017-07-17) (K. Wang et al. 2010) annotation available from the FUMA interface was used to associate SNPs to functional elements in the genome. 61 uSNPs were not in ANNOVAR. Functional annotation of those 61 SNPs was performed based on intersection with protein-coding gene coordinates. SNPs not overlapping protein-coding gene annotations were classified as intergenic. We also identified SNPs overlapping protein-coding promoters, defined as regions spanning 5 kb upstream and 1 kb downstream of protein-coding TSSs from Ensembl (v109, (Cunningham et al. 2022)).

To annotate the regulatory potential of uSNPs, we combined genomic annotations including histone modifications and chromatin accessibility across relevant human tissues. We downloaded ENCODE (v3 2021, hg38 (Dunham et al. 2012)) annotations of 1,063,878 cis-regulatory elements across 1,518 cell types and tissues. Using bedtools intersect (-wao), we annotated uSNP mapping to predicted promoters and enhancers (proximal or distal) or CTCF sites in any human tissue or cell type. We used ReMap 2022 to identify uSNPs overlapping transcription factor binding peaks (Hammal et al. 2022). We downloaded non-redundant peaks for all 1,210 TF datasets present in ReMap across all cell types and tissues. The annotation of single SNPs was performed with bedtools intersect and a minimum overlap of 1bp. Variants overlapping peaks in uterine cell lines (myofibroblasts, endometrial stromal cells, decidualized stromal cells), healthy uterine tissues (uterus, endometrium, myometrial) or diseased uterine samples (leiomyomas, primary endometrial cancer) were annotated as ‘REMAP peak uterus’, in contrast to uSNPs overlapping any TF peak in REMAP (’ REMAP peak all’). We also identified uSNPs annotated as eQTLs by exact matching in at least one tissue in the GTEx (v10) database. Because GTEx contains only a small number of uterine eQTLs, we elected to retain all eQTLs as potentially relevant. Finally, we used the Ananastra database (Abramov et al. 2021) to annotate significant uSNPs potentially disrupting a transcription factor binding site, using the European linkage disequilibrium map and a FDR of 0.05.

### Genomic 3D contacts in endometrial stromal and epithelial cells

Promoter capture Hi-C (PCHi-C) data on endometrial decidualised stromal cells derived from a full-term placenta was retrieved from (Sakabe et al. 2020). Bed files were converted to hg38 coordinates using the UCSC Liftover tool. uSNP coordinates were intersected with either PCHi-C baits or significant contacts.

Hi-C datasets on endometrial epithelial organoids from two separate donors were retrieved from (Hewitt et al. 2022). We removed contacts that overlapped ENCODE blacklisted regions using ‘`pairToBed --type neither`’. We then selected consensus contacts found in both donors, resulting in a list of 3,873 endometrial epithelial 3D genomic contacts. Those contacts were intersected with the coordinates of human protein-coding promoters, defined as a −1kb/+5kb window around gene TSSs, with (`pairToBed --type either`).

uSNPs overlapping PCHi-C or Hi-C contacts in stromal and/or epithelial cells were associated to putative target genes in the corresponding Hi-C contact, *i.e.* one or two genes depending on the type of contact (CRE-promoter, or promoter-promoter), resulting in a list of 246 and 61 uSNPs in endometrial stromal and epithelial contacts respectively.

### Reanalysis of single-nuclei RNA and ATAC-seq of healthy endometrial cells

Single-nuclei RNA-seq data of endometrial tissue from the HECA atlas (Marečková et al. 2023) was re-analysed using Seurat (v5). The HECA atlas was reduced to samples obtained from healthy donors without hormonal treatment. The resulting dataset was re-clustered and cluster annotation was homogenised based on the endometrial cell type annotation available in HECA. Nuclei from the uterine cervix (Epithelial MUC5B: n=1000; and Stromal HOXA13: n= 1471) were removed from the resulting atlas to only consider cells of endometrial origin. The resulting annotation of endometrial cell populations is presented in **Figure S4**. Pseudobulk transcriptomes were generated for each endometrial cell population using aggregate.Matrix, and scaled to the raw library size of each pseudobulk using edgeR calcNormFactors. Resulting pseudobulks were normalised to log2CPM (counts per million) and used to map the expression of uSNPs target genes across endometrial cell types.

Single-nuclei ATAC-seq data from healthy human endometrium samples was retrieved from Vrljicak et al. (2023). Data was reanalysed using the same parameters as the original analysis to filter low quality nuclei and call ATAC-seq peaks (nCount_peaks > 3000; nCount_peaks < 30000; nucleosome signal > 4; TSS enrichment > 3). To infer cell types, we combined the original cell lineage annotation from Vrljicak et al. with snRNA-seq annotation predictions using the HECA atlas (after cell type harmonisation, see above). To perform this step, gene scores were predicted for snATAC-seq nuclei using *GeneActivity*() from the Signac package (Stuart et al. 2021). These gene scores were then correlated with gene expression levels in the snRNA-seq data to transfer cell annotations using *FindTransferAnchors* and *TransferData* with default parameters. The resulting annotation is shown in **Figure S6**. Peak calling was performed by cell type using Signac with default parameters. uSNPs were then overlapped with accessible peaks in each cell type.

### Tissue-specific enrichment of uSNP target genes with FUMA

Putative uSNP target genes were passed to the FUMA GENE2FUNC interface to compute expression enrichment across 30 tissues from GTEx. Differentially expressed genes (DEG) sets in FUMA are defined using a two-sided t-test per tissue against all other tissues, with a Bonferroni corrected p-value <0.05 and a minimal absolute fold-change cut-off. The fold-enrichment in the present study represents the hypergeometric enrichment of uSNP target genes in tissue-specific DEGs.

### Gene Ontology analysis

Gene Ontology (GO) enrichment analysis was performed with the clusterProfiler package (v4.16) (T. Wu et al. 2021) in R by comparing the 227 uSNP target genes to the full transcriptome. Gene sets from the "Biological processes" category were used for gene set enrichment analysis. Enrichment was considered significant for p < 0.05 after Benjamini-Hochberg correction for multiple testing. Significant biological processes were clustered using the Ward distance method to group higher-order terms based on shared genes. The comprehensive results of the GO analysis are presented in **Table S8**.

### Analysis of uSNP derived allele effect sizes

The 1000 Genomes Phase 3 vcf files (Auton et al. 2015) were used as input to retrieve the ancestral and derived alleles for every SNP, determined using the six primates (human, chimpanzee, gorilla, orangutan, macaque, marmoset) EPO alignments from Ensembl. We restricted the list of 235 lead uSNPs to 211 bi-allelic lead uSNPs, for which an ancestral and a derived allele can be identified. If the alternate allele in the reference dataset corresponded to the ancestral allele, we switched effect size direction to reflect the impact of the derived allele on the trait. We then performed unsupervised K-means clustering on derived allele effect sizes across traits. The optimal number of clusters was identified using the average silhouette width criterion. To explore how derived alleles contribute to risk increase or decrease across different uterine disorders, effect sizes of the derived alleles were normalized across traits by computing the ratio of the effect size (β) to its standard error (SE_β). We then correlated derived allele effect sizes in each cluster across uterine disorders using Spearman’s rank correlation. Correlation p-values were adjusted using BH correction.

### Construction of control SNP datasets

Vcf files from 1000 Genomes Phase 3 were used as reference for all evolutionary analyses. SNPs in the 1000 Genome Phase 3 panels were further annotated with their minor allele frequencies for all 1000 Genomes populations, local gene densities, GERP scores and recombination rates. Minor allele frequencies (MAF) were computed using vcftools (--freq) and filtered to keep only dimorphic sites for each population independently. Local gene densities were calculated by binning the genome into 1 Mb blocks to generate pseudo-independent regions of the genome, and counting all annotated TSS within the 1 Mb region, including lncRNAs and pseudogenes. GERP scores were obtained from Ensembl, calculated from the 91-mammal whole genome alignments. Recombination rates from the 1000 Genomes panels were downloaded from UCSC (http://hgdownload.soe.ucsc.edu/gbdb/hg38/recombRate/recomb1000GAvg.bw).

For each lead uSNP present in the 1000 Genomes panels, control SNPs were defined as SNPs matched for MAF, recombination rate and gene density. To control for recombination rate and gene density, SNPs were ranked and divided into 20 bins of equal size. Control SNPs meet the following criteria: (1) with a MAF within 0.025 of their corresponding lead uSNP based on allele frequencies in the CEU population; (2) within the same recombination rate bin; (3) within the same gene density bin; and (4) excluding SNPs in LD (r^2^ > 0.2) with their corresponding lead uSNP. For each lead uSNP, the resulting list of matching control SNPs was sampled 1000 times at random with replacement to create 1000 control sets per lead uSNP. Controls SNPs across lead uSNPs were further combined to form 1000 sets of control SNPs and used to compare F_ST_ distributions. Out of 241 lead uSNPs, 50 either were not present in 1000 Genomes panels or could not be assigned appropriate matching control sets, and were excluded from the comparison.

### F_ST_ score comparisons

F_ST_ scores (Weir et Cockerham 1984) were computed from the 1000 Genomes Phase 3 vcf files using the software selink (https://github.com/h-e-g/selink). Pairwise F_ST_ scores were computed between three 1000 Genomes populations of European, East Asian and African ancestry (respectively CEU, CHB and YRI). To test for polygenic selection signatures, we compared the distributions of F_ST_ scores between lead uSNPs and matched control sets (see above) using Wilcoxon rank sum tests. P-values were BH-corrected for multiple testing across population pairs (CEU vs. CHB, CEU vs. YRI and CHB vs. YRI).

For each pair of populations, a directional signed F_ST_ score (positive or negative) defined as previously proposed in (Quach et al. 2016), to orient whether the derived allele is more or less frequent with respect to a reference population. Directional F_ST_ scores were used to identify lead uSNPs with outlier F_ST_ scores and under putative selection in either European, East-Asian or African populations. Outlier F_ST_ SNPs were defined as the top 5% genome-wide F_ST_ signals for each population comparison. To test for enrichment in outlier F_ST_ SNPs, we compared the fraction of lead uSNPs in the top 5% F_ST_ values to the fraction of controls across all 1000 control sets and derived an empirical p-value. Resulting p-values were BH-corrected for multiple testing across population pairs (CEU vs. CHB, CEU vs. YRI and CHB vs. YRI). We also tested for an overall enrichment of lead uSNPs in the top 5% F_ST_ signals using a two-sided binomial test (probability of finding a uSNPs in the top 5% in at least one F_ST_ comparison = 1-0.95^3^).

### iHS score comparisons

iHS scores were obtained from Johnson et Voight (2018). We compared distributions of absolute iHS scores between lead uSNPs and matched control sets using Wilcoxon rank sum tests. Out of 241 lead uSNPs, only 163 lead uSNPs had an available iHS value in at least one population.

### Code availability

All original code produced during this project is accessible at https://gitlab.pasteur.fr/euliorzo/popgen. JASS is available at https://statistical-genetics.pages.pasteur.fr/jass/index.html.

## Licences

Figures were produced with BioRender under licence to Institut Pasteur.

## Acknowledgments

We thank Jan Brosens, Pavle Vrljicak and Mireia Taus-Nebot (University of Warwick) for sharing analyzed snATAC-seq datasets from their previous study.

## Funding

This project was supported by Institut Pasteur (G5 package), Centre National de la Recherche Scientifique (CNRS UMR 3525), Institut National de la Santé et de la Recherche Médicale (INSERM UA12), and the Inception program (Investissement d’Avenir grant ANR-16-CONV-0005). EL is supported by a PhD fellowship from Université Paris Cité and a grant from the Fondation pour la Recherche Médicale (grant agreement FDT202504020295).

## Authors contributions

EL and CB conceived and designed the project and analyses. EL performed all analyses, with contributions from HJ and HA for multi-trait GWAS analyses, MD for functional genomics analyses, and GL for evolutionary analyses. EL designed all figures. EL and CB wrote the manuscript with input from all authors. All authors approved the manuscript.

## Competing interests

The authors declare no competing interests.

## Notes

### Competing Interest Statement

The authors have declared no competing interest.

## References

Abramov, Sergey, Alexandr Boytsov, Daria Bykova, et al. 2021. « Landscape of Allele-Specific Transcription Factor Binding in the Human Genome ». Nature Communications 12 (1): 1. 10.1038/s41467-021-23007-0.

Auton, Adam, Gonçalo R. Abecasis, David M. Altshuler, et al. 2015. « A Global Reference for Human Genetic Variation ». Nature 526 (7571): 68–74. 10.1038/nature15393.

Benonisdottir, Stefania, Vincent J. Straub, Augustine Kong, et Melinda C. Mills. 2024. « Genetics of Female and Male Reproductive Traits and Their Relationship with Health, Longevity and Consequences for Offspring ». Nature Aging 4 (12): 1745–59. 10.1038/s43587-024-00733-w.

Berg, Jeremy J., Arbel Harpak, Nasa Sinnott-Armstrong, et al. 2019. « Reduced Signal for Polygenic Adaptation of Height in UK Biobank ». eLife 8 (mars): e39725. 10.7554/eLife.39725.

Bourdon, Mathilde, Pietro Santulli, Louis Marcellin, et al. 2021. « Adenomyosis: An update regarding its diagnosis and clinical features ». Journal of Gynecology Obstetrics and Human Reproduction 50 (10): 102228. 10.1016/j.jogoh.2021.102228.

Bulik-Sullivan, Brendan, Hilary K. Finucane, Verneri Anttila, et al. 2015. « An Atlas of Genetic Correlations across Human Diseases and Traits ». Nature Genetics 47 (11): 1236–41. 10.1038/ng.3406.

Bulik-Sullivan, Brendan K., Po-Ru Loh, Hilary K. Finucane, et al. 2015. « LD Score Regression Distinguishes Confounding from Polygenicity in Genome-Wide Association Studies ». Nature Genetics 47 (3): 291–95. 10.1038/ng.3211.

Bulun, Serdar E., Bahar D. Yilmaz, Christia Sison, et al. 2019. « Endometriosis ». Endocrine Reviews 40 (4): 1048–79. 10.1210/er.2018-00242.

Bulun, Serdar E., Ping Yin, JianJun Wei, et al. 2025. « Uterine Fibroids ». Physiological Reviews, publication en ligne anticipée, juin 13. Rockville, MD. 10.1152/physrev.00010.2024.

Buyukcelebi, Kadir, Alexander J. Duval, Fatih Abdula, Hoda Elkafas, Fidan Seker-Polat, et Mazhar Adli. 2024. « Integrating Leiomyoma Genetics, Epigenomics, and Single-Cell Transcriptomics Reveals Causal Genetic Variants, Genes, and Cell Types ». Nature Communications 15 (1): 1169. 10.1038/s41467-024-45382-0.

Cardoso, Jéssica Vilarinho, Jamila Alessandra Perini, Daniel Escorsim Machado, Ricardo Pinto, et Rui Medeiros. 2020. « Systematic review of genome-wide association studies on susceptibility to endometriosis ». European Journal of Obstetrics & Gynecology and Reproductive Biology 255 (décembre): 74–82. 10.1016/j.ejogrb.2020.10.017.

Chen, Piaopiao, et Jianzhi Zhang. 2020. « Antagonistic Pleiotropy Conceals Molecular Adaptations in Changing Environments ». Nature Ecology & Evolution 4 (3): 461–69. 10.1038/s41559-020-1107-8.

Choi, Eun Ji, Seong Beom Cho, Sa Ra Lee, et al. 2017. « Comorbidity of gynecological and non-gynecological diseases with adenomyosis and endometriosis ». Obstetrics & Gynecology Science 60 (6): 579–86. 10.5468/ogs.2017.60.6.579.

Choin, Jeremy, Javier Mendoza-Revilla, Lara R. Arauna, et al. 2021. « Genomic Insights into Population History and Biological Adaptation in Oceania ». Nature 592 (7855): 7855. 10.1038/s41586-021-03236-5.

Constantinescu, Andrei-Emil, Ruth E. Mitchell, Jie Zheng, et al. 2022. « A framework for research into continental ancestry groups of the UK Biobank ». Human Genomics 16 (1): 3. 10.1186/s40246-022-00380-5.

Cunningham, Fiona, James E. Allen, Jamie Allen, et al. 2022. « Ensembl 2022 ». Nucleic Acids Research 50 (D1): D988–95. 10.1093/nar/gkab1049.

Dong, Di, Zhe Zhou, Minglu Zhu, et al. 2025. « STN1 Facilitates Metastasis by Promoting Transcription of EMT-Activator ZEB1 in Pancreatic Cancer ». Nature Communications 16 (1): 7815. 10.1038/s41467-025-63083-0.

Dunham, Ian, Anshul Kundaje, Shelley F. Aldred, et al. 2012. « An Integrated Encyclopedia of DNA Elements in the Human Genome ». Nature 489 (7414): 57–74. 10.1038/nature11247.

Fiore, Alexander, Maìra Casalechi, Laura Sichenze, et al. 2025. « Co-Occurrence of Endometriosis and Uterine Fibroids: A Systematic Review and Meta-Analysis ». eClinicalMedicine 89 (novembre). 10.1016/j.eclinm.2025.103510.

Fraser, Ian S., Hilary O. D. Critchley, Michael Broder, et Malcolm G. Munro. 2011. « The FIGO Recommendations on Terminologies and Definitions for Normal and Abnormal Uterine Bleeding ». Seminars in Reproductive Medicine 29 (5): 383–90. 10.1055/s-0031-1287662.

Gallagher, C. S., N. Mäkinen, H. R. Harris, et al. 2019. « Genome-Wide Association and Epidemiological Analyses Reveal Common Genetic Origins between Uterine Leiomyomata and Endometriosis ». Nature Communications 10 (1): 4857. 10.1038/s41467-019-12536-4.

Gao, J., S. Zeng, B. L. Sun, H. M. Fan, et L. H. Han. 1987. « Menstrual Blood Loss and Hematologic Indices in Healthy Chinese Women ». The Journal of Reproductive Medicine 32 (11): 822–26.

Giuliani, Emma, Sawsan As-Sanie, et Erica E. Marsh. 2020. « Epidemiology and Management of Uterine Fibroids ». International Journal of Gynecology & Obstetrics 149 (1): 3–9. 10.1002/ijgo.13102.

GTEx Consortium. 2017. « Genetic Effects on Gene Expression across Human Tissues ». Nature 550 (7675): 204–13. 10.1038/nature24277.

Guare, Lindsay A., Jagyashila Das, Lannawill Caruth, et al. 2025. « Expanding the Genetic Landscape of Endometriosis: Integrative -Omics Analyses Implicate Key Genes and Pathways in a Multi-Ancestry Study of over One Million Women ». Prépublication, medRxiv, novembre 14. 10.1101/2024.11.26.24316723.

Hämälä, Tuomas, Amanda J. Gorton, David A. Moeller, et Peter Tiffin. 2020. « Pleiotropy Facilitates Local Adaptation to Distant Optima in Common Ragweed (Ambrosia Artemisiifolia) ». PLOS Genetics 16 (3): e1008707. 10.1371/journal.pgen.1008707.

Hammal, Fayrouz, Pierre de Langen, Aurélie Bergon, Fabrice Lopez, et Benoit Ballester. 2022. « ReMap 2022: A Database of Human, Mouse, Drosophila and Arabidopsis Regulatory Regions from an Integrative Analysis of DNA-Binding Sequencing Experiments ». Nucleic Acids Research 50 (D1): D316–25. 10.1093/nar/gkab996.

Hewitt, Sylvia C., San-pin Wu, Tianyuan Wang, et al. 2022. « Progesterone Signaling in Endometrial Epithelial Organoids ». Cells 11 (11). 10.3390/cells11111760.

Holsinger, Kent E., et Bruce S. Weir. 2009. « Genetics in Geographically Structured Populations: Defining, Estimating and Interpreting FST ». Nature Reviews Genetics 10 (9): 9. 10.1038/nrg2611.

Johnson, Kelsey Elizabeth, et Benjamin F. Voight. 2018. « Patterns of Shared Signatures of Recent Positive Selection across Human Populations ». Nature Ecology & Evolution 2 (4): 713–20. 10.1038/s41559-018-0478-6.

Julienne, Hanna, Vincent Laville, Zachary R. McCaw, et al. 2021. « Multitrait GWAS to Connect Disease Variants and Biological Mechanisms ». PLOS Genetics 17 (8): e1009713. 10.1371/journal.pgen.1009713.

Julienne, Hanna, Pierre Lechat, Vincent Guillemot, et al. 2020. « JASS: command line and web interface for the joint analysis of GWAS results ». NAR Genomics and Bioinformatics 2 (1): lqaa003. 10.1093/nargab/lqaa003.

Karczewski, Konrad J., Rahul Gupta, Masahiro Kanai, et al. 2025. « Pan-UK Biobank Genome-Wide Association Analyses Enhance Discovery and Resolution of Ancestry-Enriched Effects ». Nature Genetics 57 (10): 2408–17. 10.1038/s41588-025-02335-7.

Kim, Jeewoo, Ariel Williams, Hannah Noh, et al. 2024. « 430 Genome-Wide Meta-Analysis Identifies Novel Risk Loci for Uterine Fibroids across Multiple Ancestry Groups ». Journal of Clinical and Translational Science 8 (s1): 128–29. 10.1017/cts.2024.372.

Koller, Dora, Jun He, Solveig Løkhammer, et al. 2025. « Multi-Ancestry Genome-Wide Association Study of Endometriosis and Its Clinical Manifestations in ∼1.4 Million Women: Translating Gene Discovery into Pathogenic Mechanisms and Therapeutic Targets ». Prépublication, medRxiv, septembre 5. 10.1101/2025.09.03.25335012.

Kun, Eucharist, Emily M. Javan, Olivia Smith, et al. 2023. « The Genetic Architecture and Evolution of the Human Skeletal Form ». Science 381 (6655): eadf8009. 10.1126/science.adf8009.

Kurki, Mitja I., Juha Karjalainen, Priit Palta, et al. 2022. FinnGen: Unique Genetic Insights from Combining Isolated Population and National Health Register Data. Preprint. Genetic and Genomic Medicine. 10.1101/2022.03.03.22271360.

Kurki, Mitja I., Juha Karjalainen, Priit Palta, et al. 2023. « FinnGen Provides Genetic Insights from a Well-Phenotyped Isolated Population ». Nature 613 (7944): 508–18. 10.1038/s41586-022-05473-8.

Kyama, Cleophas M., Jason M. Mwenda, James Machoki, et al. 2007. « Endometriosis in African Women ». Women’s Health 3 (5): 629–35. 10.2217/17455057.3.5.629.

La Vecchia, I., A. Fiore, N. Salmeri, et al. 2025. « O-199 The association between endometriosis and endometrial polyps: a systematic review and meta-analysis ». Human Reproduction 40 (Supplement_1): deaf097.199. 10.1093/humrep/deaf097.199.

Lahdenperä, Mirkka, Virpi Lummaa, Samuli Helle, Marc Tremblay, et Andrew F. Russell. 2004. « Fitness Benefits of Prolonged Post-Reproductive Lifespan in Women ». Nature 428 (6979): 178–81. 10.1038/nature02367.

Laval, Guillaume, Etienne Patin, Pierre Boutillier, et Lluis Quintana-Murci. 2021. « Sporadic occurrence of recent selective sweeps from standing variation in humans as revealed by an approximate Bayesian computation approach ». Genetics 219 (4): iyab161. 10.1093/genetics/iyab161.

Levin, Michael G., Noah L. Tsao, Pankhuri Singhal, et al. 2022. « Genome-Wide Association and Multi-Trait Analyses Characterize the Common Genetic Architecture of Heart Failure ». Nature Communications 13 (1): 6914. 10.1038/s41467-022-34216-6.

Long, Erping, et Jianzhi Zhang. 2023. « Evidence for the role of selection for reproductively advantageous alleles in human aging ». Science Advances 9 (49): eadh4990. 10.1126/sciadv.adh4990.

Lucas, Emma S., Nigel P. Dyer, Katherine Fishwick, Sascha Ott, et Jan J. Brosens. 2016. « Success after Failure: The Role of Endometrial Stem Cells in Recurrent Miscarriage ». Reproduction. Reproduction 152 (5): R159–66. 10.1530/REP-16-0306.

Makker, Vicky, Helen MacKay, Isabelle Ray-Coquard, et al. 2021. « Endometrial Cancer ». Nature Reviews Disease Primers 7 (1): 88. 10.1038/s41572-021-00324-8.

Marečková, Magda, Luz Garcia-Alonso, Marie Moullet, et al. 2023. An Integrated Single-Cell Reference Atlas of the Human Endometrium. Preprint. Genomics. 10.1101/2023.11.03.564728.

Masuda, Tatsuo, Siew-Kee Low, Masato Akiyama, et al. 2020. « GWAS of Five Gynecologic Diseases and Cross-Trait Analysis in Japanese ». European Journal of Human Genetics 28 (1): 95–107. 10.1038/s41431-019-0495-1.

McGrath, Isabelle M., Grant W. Montgomery, Sally Mortlock, et International Endometriosis Genetics Consortium. 2023. « Genomic Characterisation of the Overlap of Endometriosis with 76 Comorbidities Identifies Pleiotropic and Causal Mechanisms Underlying Disease Risk ». Human Genetics 142 (9): 1345–60. 10.1007/s00439-023-02582-w.

Mecha, Ezekiel O., Joseph N. Njagi, Roselydiah N. Makunja, Charles O. A. Omwandho, Philippa T. K. Saunders, et Andrew W. Horne. 2022. « Endometriosis among African Women ». Reproduction and Fertility. Reproduction and Fertility 3 (3): C40–43. 10.1530/RAF-22-0040.

Mortlock, Sally, Rosario I. Corona, Pik Fang Kho, et al. 2022. « A Multi-Level Investigation of the Genetic Relationship between Endometriosis and Ovarian Cancer Histotypes ». Cell Reports Medicine 3 (3): 100542. 10.1016/j.xcrm.2022.100542.

Munro, Malcolm G. 2019. « Uterine polyps, adenomyosis, leiomyomas, and endometrial receptivity ». Fertility and Sterility 111 (4): 629–40. 10.1016/j.fertnstert.2019.02.008.

Muter, Joanne, Chow-Seng Kong, et Jan J. Brosens. 2021. « The Role of Decidual Subpopulations in Implantation, Menstruation and Miscarriage ». Frontiers in Reproductive Health 3. https://www.frontiersin.org/articles/10.3389/frph.2021.804921.

Nguyen, Dinh Duc, Eugene Kim, Nhat Thong Le, et al. 2023. « Deficiency in mammalian STN1 promotes colon cancer development via inhibiting DNA repair ». Science Advances 9 (19): eadd8023. 10.1126/sciadv.add8023.

Oehler, M. K., et M. C. P. Rees. 2003. « Menorrhagia: An Update ». Acta Obstetricia et Gynecologica Scandinavica 82 (5): 405–22. 10.1034/j.1600-0412.2003.00097.x.

Painter, Jodie N., Tracy A. O’Mara, Andrew P. Morris, et al. 2018. « Genetic overlap between endometriosis and endometrial cancer: evidence from cross-disease genetic correlation and GWAS meta-analyses ». Cancer Medicine 7 (5): 1978–87. 10.1002/cam4.1445.

Pavlicev, Mihaela, Caitlin E. McDonough-Goldstein, Andreja Moset Zupan, et al. 2022. A SNP Affects Wnt4 Expression in Endometrial Stroma, with Antagonistic Implications for Pregnancy, Endometriosis and Reproductive Cancers. Preprint. Genetics. 10.1101/2022.10.25.513653.

Pavlicev, Mihaela, et Günter P. Wagner. 2012. « A model of developmental evolution: selection, pleiotropy and compensation ». Trends in Ecology & Evolution 27 (6): 316–22. 10.1016/j.tree.2012.01.016.

Pritchard, Jonathan K., et Anna Di Rienzo. 2010. « Adaptation – Not by Sweeps Alone ». Nature Reviews Genetics 11 (10): 665–67. 10.1038/nrg2880.

Pritchard, Jonathan K., Joseph K. Pickrell, et Graham Coop. 2010. « The Genetics of Human Adaptation: Hard Sweeps, Soft Sweeps, and Polygenic Adaptation ». Current Biology 20 (4): R208–15. 10.1016/j.cub.2009.11.055.

Pujol Gualdo, Natàlia, Jelisaveta Džigurski, Valentina Rukins, et al. 2025. « Atlas of Genetic and Phenotypic Associations across 42 Female Reproductive Health Diagnoses ». Nature Medicine 31 (5): 1626–34. 10.1038/s41591-025-03543-8.

Purcell, Shaun, Benjamin Neale, Kathe Todd-Brown, et al. 2007. « PLINK: A Tool Set for Whole-Genome Association and Population-Based Linkage Analyses ». American Journal of Human Genetics 81 (3): 559–75. 10.1086/519795.

Quach, Hélène, Maxime Rotival, Julien Pothlichet, et al. 2016. « Genetic Adaptation and Neandertal Admixture Shaped the Immune System of Human Populations ». Cell 167 (3): 643–656.e17. 10.1016/j.cell.2016.09.024.

Rafnar, Thorunn, Bjarni Gunnarsson, Olafur A. Stefansson, et al. 2018a. « Variants Associating with Uterine Leiomyoma Highlight Genetic Background Shared by Various Cancers and Hormone-Related Traits ». Nature Communications 9 (1): 3636. 10.1038/s41467-018-05428-6.

Rafnar, Thorunn, Bjarni Gunnarsson, Olafur A. Stefansson, et al. 2018b. « Variants Associating with Uterine Leiomyoma Highlight Genetic Background Shared by Various Cancers and Hormone-Related Traits ». Nature Communications 9 (1): 3636. 10.1038/s41467-018-05428-6.

Rahmioglu, Nilufer, Sally Mortlock, Marzieh Ghiasi, et al. 2023. « The Genetic Basis of Endometriosis and Comorbidity with Other Pain and Inflammatory Conditions ». Nature Genetics 55 (3): 3. 10.1038/s41588-023-01323-z.

Rodríguez, Juan Antonio, Urko M. Marigorta, David A. Hughes, Nino Spataro, Elena Bosch, et Arcadi Navarro. 2017a. « Antagonistic Pleiotropy and Mutation Accumulation Influence Human Senescence and Disease ». Nature Ecology & Evolution 1 (3): 0055. 10.1038/s41559-016-0055.

Rodríguez, Juan Antonio, Urko M. Marigorta, David A. Hughes, Nino Spataro, Elena Bosch, et Arcadi Navarro. 2017b. « Antagonistic Pleiotropy and Mutation Accumulation Influence Human Senescence and Disease ». Nature Ecology & Evolution 1 (3): 0055. 10.1038/s41559-016-0055.

Rose, Michael R. 1982. « Antagonistic Pleiotropy, Dominance, and Genetic Variation ». Heredity 48 (1): 63–78. 10.1038/hdy.1982.7.

Sakabe, Noboru J., Ivy Aneas, Nicholas Knoblauch, et al. 2020. « Transcriptome and regulatory maps of decidua-derived stromal cells inform gene discovery in preterm birth ». Science Advances 6 (49): eabc8696. 10.1126/sciadv.abc8696.

Sapkota, Yadav, Valgerdur Steinthorsdottir, Andrew P. Morris, et al. 2017a. « Meta-Analysis Identifies Five Novel Loci Associated with Endometriosis Highlighting Key Genes Involved in Hormone Metabolism ». Nature Communications 8 (1): 15539. 10.1038/ncomms15539.

Sapkota, Yadav, Valgerdur Steinthorsdottir, Andrew P. Morris, et al. 2017b. « Meta-Analysis Identifies Five Novel Loci Associated with Endometriosis Highlighting Key Genes Involved in Hormone Metabolism ». Nature Communications 8 (1): 15539. 10.1038/ncomms15539.

Sheng, Xin, Yuting Guan, Ziyuan Ma, et al. 2021. « Mapping the Genetic Architecture of Human Traits to Cell Types in the Kidney Identifies Mechanisms of Disease and Potential Treatments ». Nature Genetics 53 (9): 1322–33. 10.1038/s41588-021-00909-9.

Sheu, Jim Jinn-Chyuan, Wei-Yong Lin, Ting-Yuan Liu, et al. 2024a. « Ethnic-Specific Genetic Susceptibility Loci for Endometriosis in Taiwanese-Han Population: A Genome-Wide Association Study ». Journal of Human Genetics 69 (11): 573–83. 10.1038/s10038-024-01270-5.

Sheu, Jim Jinn-Chyuan, Wei-Yong Lin, Ting-Yuan Liu, et al. 2024b. « Ethnic-Specific Genetic Susceptibility Loci for Endometriosis in Taiwanese-Han Population: A Genome-Wide Association Study ». Journal of Human Genetics, juillet 9, 1–11. 10.1038/s10038-024-01270-5.

Sinharoy, Sheela S., Lyzberthe Chery, Madeleine Patrick, et al. 2023. « Prevalence of Heavy Menstrual Bleeding and Associations with Physical Health and Wellbeing in Low-Income and Middle-Income Countries: A Multinational Cross-Sectional Study ». The Lancet Global Health 11 (11): e1775–84. 10.1016/S2214-109X(23)00416-3.

Soliman, A. M., H. Yang, E. X. Du, S. Kelkar, et C. Winkel. 2016. « A Systematic Literature Review of Comorbidities and Symptoms among Uterine Fibroids Patients between 2000 and 2013 ». Value in Health 19 (3): A172. 10.1016/j.jval.2016.03.1446.

Song, Jiangwei, Ning Gao, Zhe Chen, et al. 2024. « Shared genetic etiology of vessel diseases: A genome-wide multi-traits association analysis ». Thrombosis Research 241 (septembre): 109102. 10.1016/j.thromres.2024.109102.

Song, Weichen, Yueqi Shi, Weidi Wang, et al. 2021. « A Selection Pressure Landscape for 870 Human Polygenic Traits ». Nature Human Behaviour 5 (12): 1731–43. 10.1038/s41562-021-01231-4.

Stephan, Wolfgang. 2016. « Signatures of Positive Selection: From Selective Sweeps at Individual Loci to Subtle Allele Frequency Changes in Polygenic Adaptation ». Molecular Ecology 25 (1): 79–88. 10.1111/mec.13288.

Stephan, Wolfgang, et Sona John. 2020. « Polygenic Adaptation in a Population of Finite Size ». Entropy 22 (8): 907. 10.3390/e22080907.

Stewart, E. A., C. L. Cookson, R. A. Gandolfo, et R. Schulze-Rath. 2017a. « Epidemiology of Uterine Fibroids: A Systematic Review ». BJOG: An International Journal of Obstetrics and Gynaecology 124 (10): 1501–12. 10.1111/1471-0528.14640.

Stewart, E. A., C. L. Cookson, R. A. Gandolfo, et R. Schulze-Rath. 2017b. « Epidemiology of Uterine Fibroids: A Systematic Review ». BJOG: An International Journal of Obstetrics and Gynaecology 124 (10): 1501–12. 10.1111/1471-0528.14640.

Stuart, Tim, Avi Srivastava, Shaista Madad, Caleb A. Lareau, et Rahul Satija. 2021. « Single-Cell Chromatin State Analysis with Signac ». Nature Methods 18 (11): 1333–41. 10.1038/s41592-021-01282-5.

The 1000 Genomes Project Consortium, Corresponding authors, Adam Auton, et al. 2015. « A Global Reference for Human Genetic Variation ». Nature 526 (7571): 68–74. 10.1038/nature15393.

Thibord, Florian, Jason Cunha, Jelisaveta Džigurski, et al. 2025. « Genome-wide meta-analysis of heavy menstrual bleeding reveals 36 risk loci ». Blood 146 (6): 745–58. 10.1182/blood.2024027382.

Venkatesh, Samvida S., Laura B. L. Wittemans, Duncan S. Palmer, et al. 2025. « Genome-Wide Analyses Identify 25 Infertility Loci and Relationships with Reproductive Traits across the Allele Frequency Spectrum ». Nature Genetics 57 (5): 1107–18. 10.1038/s41588-025-02156-8.

Voight, Benjamin F., Sridhar Kudaravalli, Xiaoquan Wen, et Jonathan K. Pritchard. 2006a. « A Map of Recent Positive Selection in the Human Genome ». PLOS Biology 4 (3): e72. 10.1371/journal.pbio.0040072.

Voight, Benjamin F., Sridhar Kudaravalli, Xiaoquan Wen, et Jonathan K. Pritchard. 2006b. « A Map of Recent Positive Selection in the Human Genome ». PLoS Biology 4 (3): e72. 10.1371/journal.pbio.0040072.

Vrljicak, Pavle, Emma S. Lucas, Maria Tryfonos, Joanne Muter, Sascha Ott, et Jan J. Brosens. 2023. « Dynamic Chromatin Remodeling in Cycling Human Endometrium at Single-Cell Level ». Cell Reports 42 (12). 10.1016/j.celrep.2023.113525.

Wang, Kai, Mingyao Li, et Hakon Hakonarson. 2010. « ANNOVAR: functional annotation of genetic variants from high-throughput sequencing data ». Nucleic Acids Research 38 (16): e164. 10.1093/nar/gkq603.

Wang, Xuan-Yu, Qiong Lyu, Yang-Yang Zhang, et al. 2025. « Shared Genetic Architecture between Metabolic Dysfunction-Associated Steatotic Liver Disease and Cardiometabolic Traits Comorbidities: A Genome-Wide Pleiotropic and Multi-Omics Study ». Metabolism and Target Organ Damage 5 (2): N/A-N/A. 10.20517/mtod.2024.129.

Wang, Xuemin, Dylan M. Glubb, et Tracy A. O’Mara. 2022. « 10 Years of GWAS Discovery in Endometrial Cancer: Aetiology, Function and Translation ». eBioMedicine 77 (mars). 10.1016/j.ebiom.2022.103895.

Weir, B. S., et C. Clark Cockerham. 1984. « Estimating F-Statistics for the Analysis of Population Structure ». Evolution 38 (6): 1358–70. 10.2307/2408641.

Wu, Dou, Zi Wang, Jingying Huang, et al. 2022. « An antagonistic pleiotropic gene regulates the reproduction and longevity tradeoff ». Proceedings of the National Academy of Sciences 119 (18): e2120311119. 10.1073/pnas.2120311119.

Wu, Tianzhi, Erqiang Hu, Shuangbin Xu, et al. 2021. « clusterProfiler 4.0: A universal enrichment tool for interpreting omics data ». The Innovation 2 (3): 100141. 10.1016/j.xinn.2021.100141.

Xiang, Yifan, Vineeta Tanwar, Parminder Singh, Lizellen La Follette, et Pankaj Kapahi. 2024. « Early Menarche and Childbirth Accelerate Aging-Related Outcomes and Age-Related Diseases: Evidence for Antagonistic Pleiotropy in Humans ». eLife 13 (novembre). 10.7554/eLife.102447.1.

Yamamoto, Ayae, Erica B. Johnstone, Michael S. Bloom, Heather G. Huddleston, et Victor Y. Fujimoto. 2017. « A Higher Prevalence of Endometriosis among Asian Women Does Not Contribute to Poorer IVF Outcomes ». Journal of Assisted Reproduction and Genetics 34 (6): 765–74. 10.1007/s10815-017-0919-1.

Ye, Jiatian, Hongling Peng, Xia Huang, et Xiaorong Qi. 2022. « The association between endometriosis and risk of endometrial cancer and breast cancer: a meta-analysis ». BMC Women’s Health 22 (novembre): 455. 10.1186/s12905-022-02028-x.

