## Supplementary Figures for "Evolution of the genetic architecture of uterine disorders"

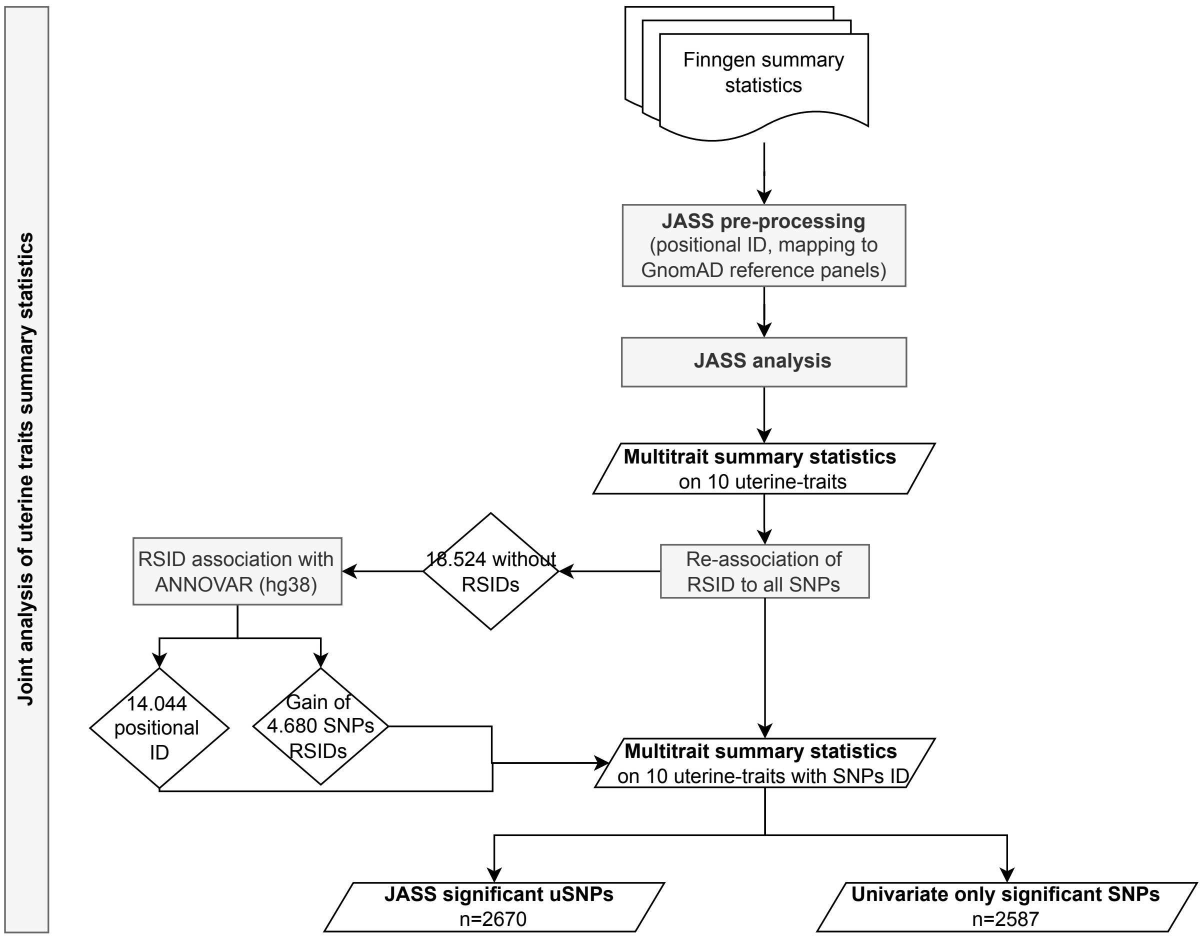

**Figure S1: Diagram of the JASS multi-trait analysis.**

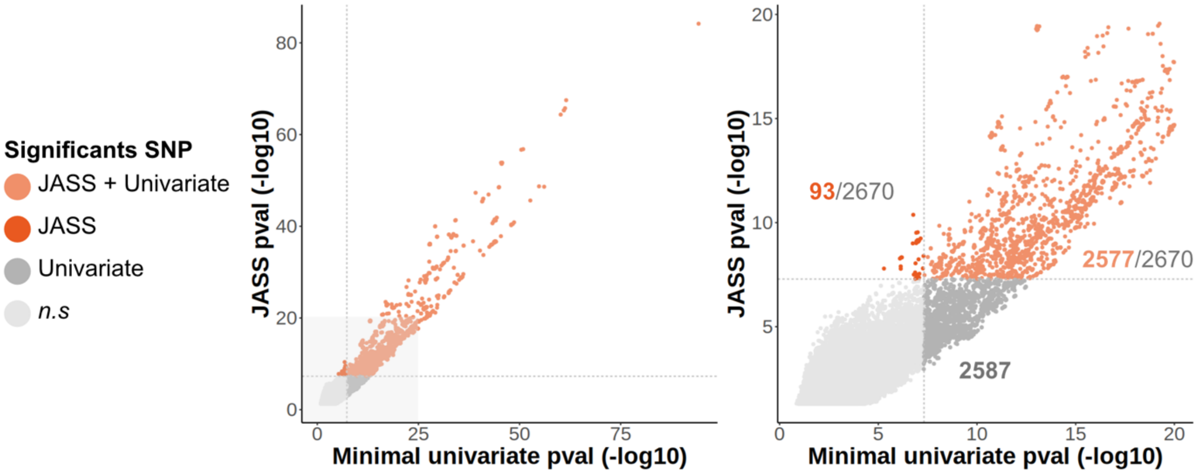

**Figure S2: Quadrant plot of significant SNPs after JASS multi-trait analysis.**

This plot shows the -log10 (p-value) of the joint association test against the lowest -log10(p-value) in single-trait GWAS for all SNPs in the FinnGen reference panel. Complete results are presented in the left panel, and a zoom around the genome-wide significance threshold is presented on the right panel.

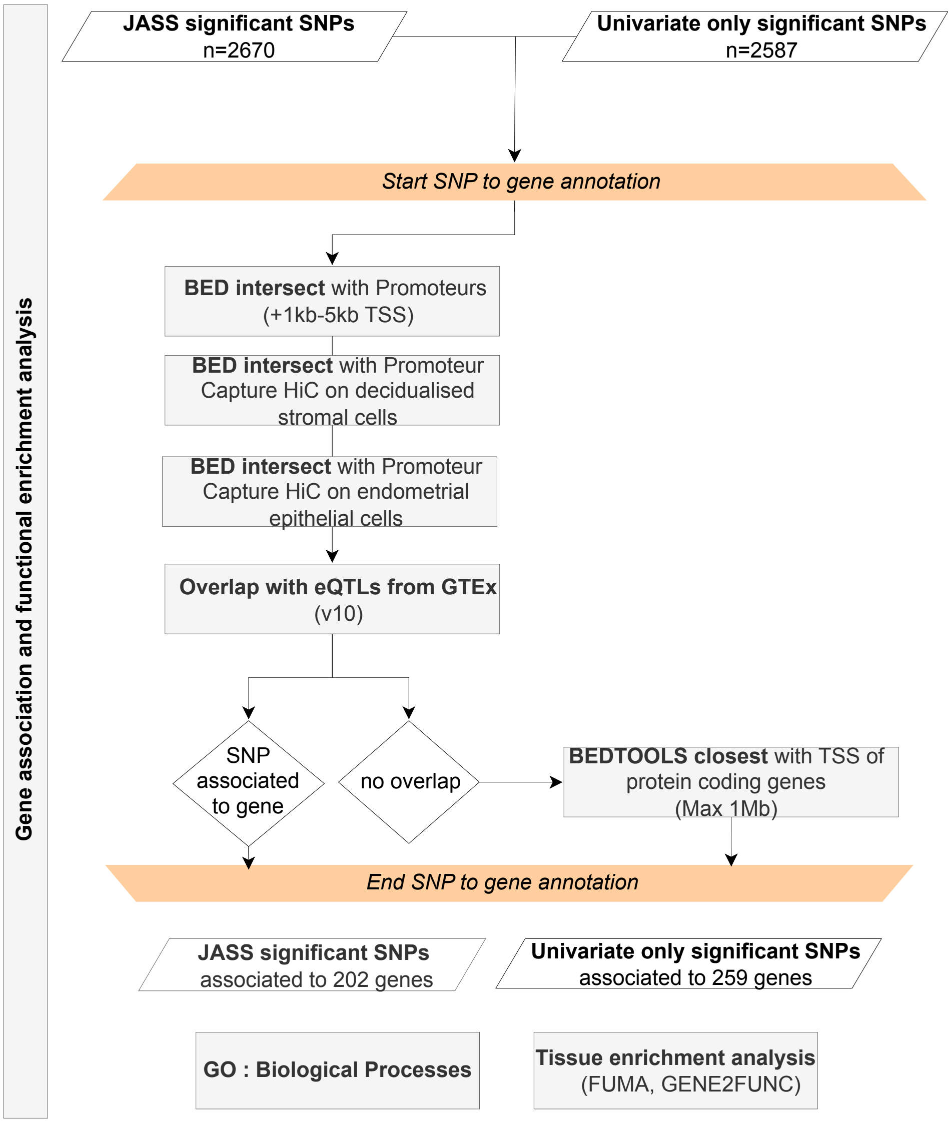

**Figure S3: Diagram of the SNP-to-Gene annotation combining functional annotation and positional annotation based on distance to gene TSS.**

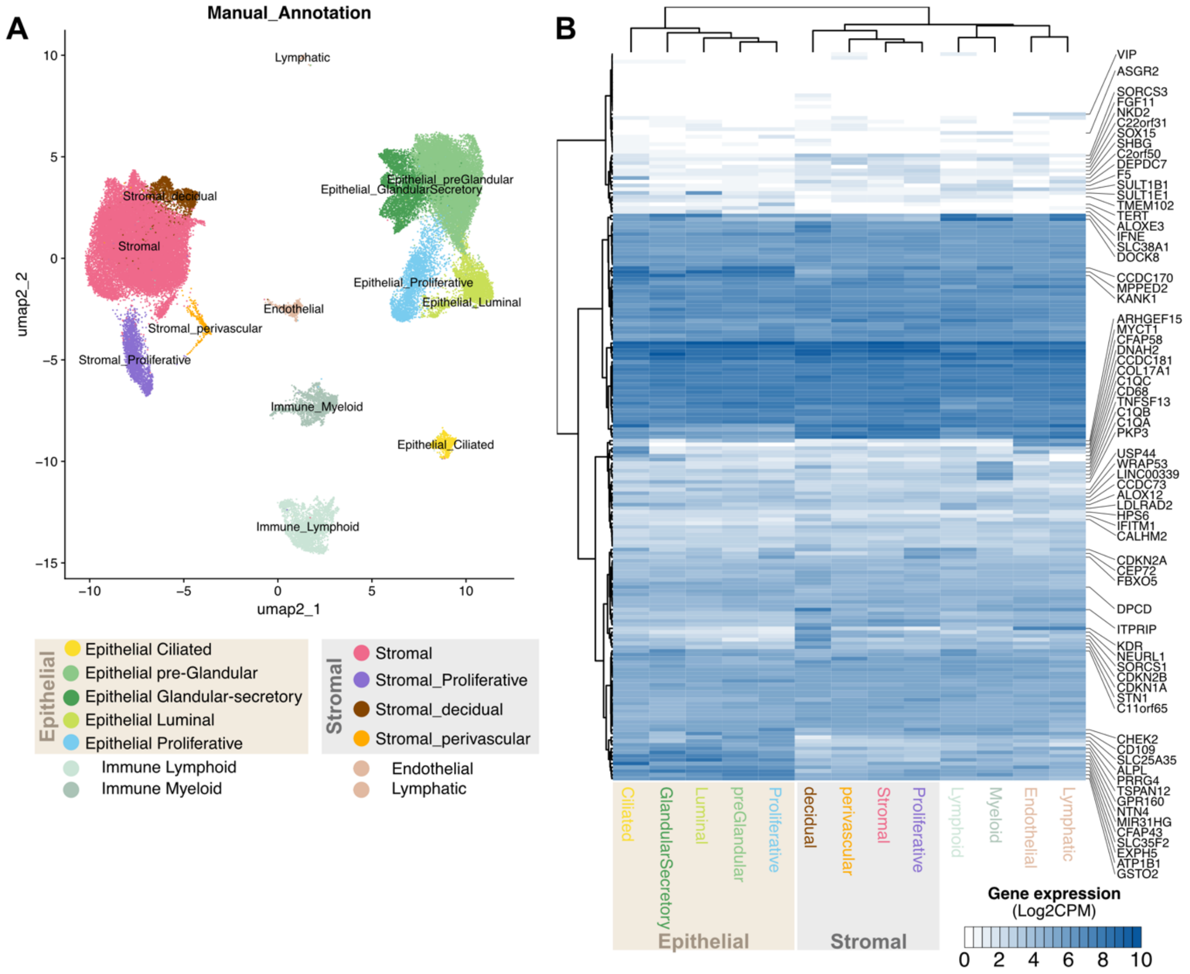

**Figure S4: Re-analysis of the healthy human endometrial single-nuclei RNA-seq atlas (HECA, (Marečková et al. 2023)).**

(A) Unifold Manifold Approximation and Projection (UMAP) of endometrial cell types from the HECA atlas after subsetting for nuclei recovered from healthy donors, without hormonal treatment.

(B) Expression heatmap for all 194 uSNP target genes (uGenes) quantified in HECA across endometrial cell types. Gene expression in normalised log2CPM counts.

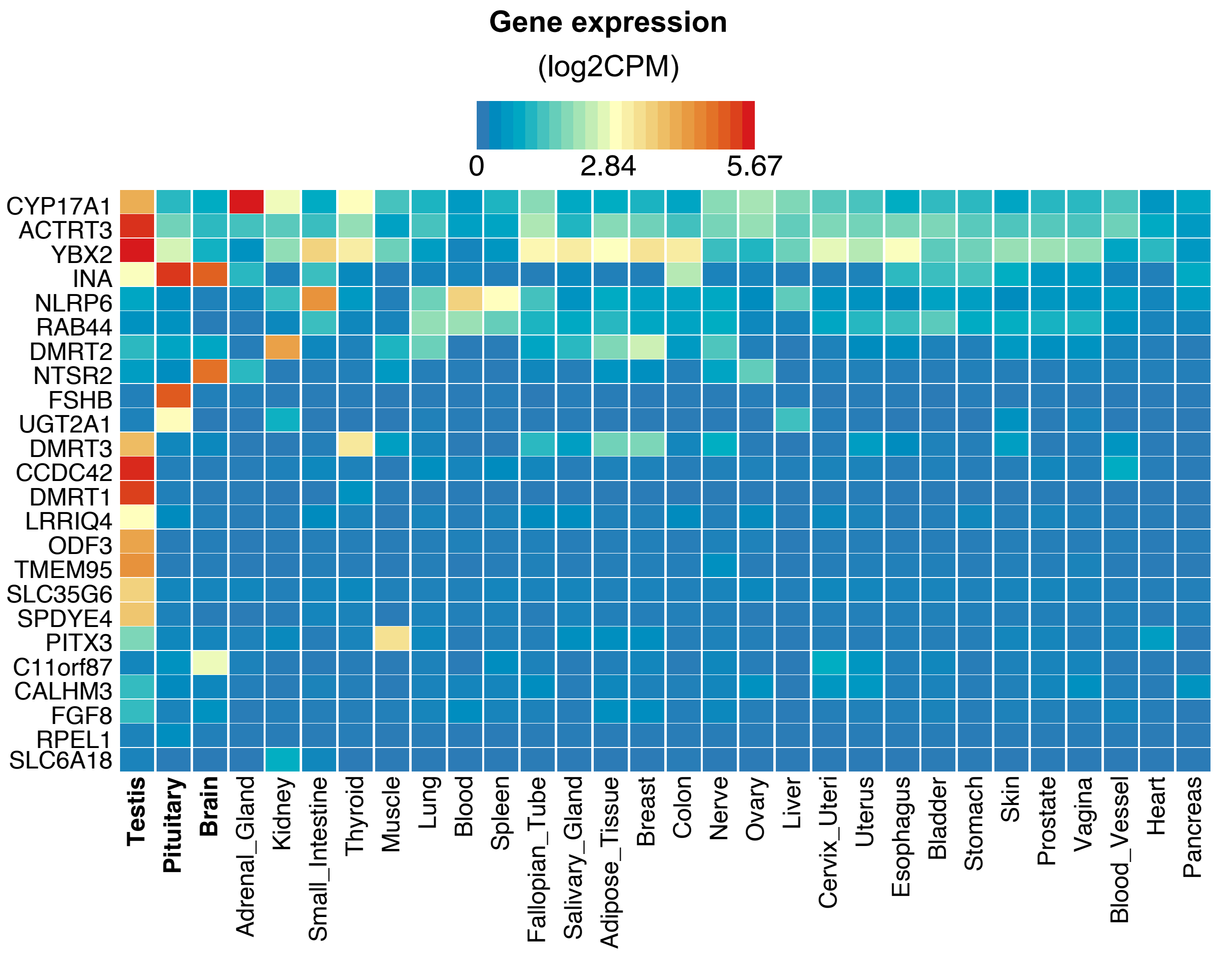

**Figure S5: Gene expression heatmap of uSNP target genes not expressed in uterine cells across other GTEx tissues.**

Heatmap of gene expression normalised (log2CPM) across GTEx tissue transcriptomes. Heatmap generated using the FUMA online toolkit.

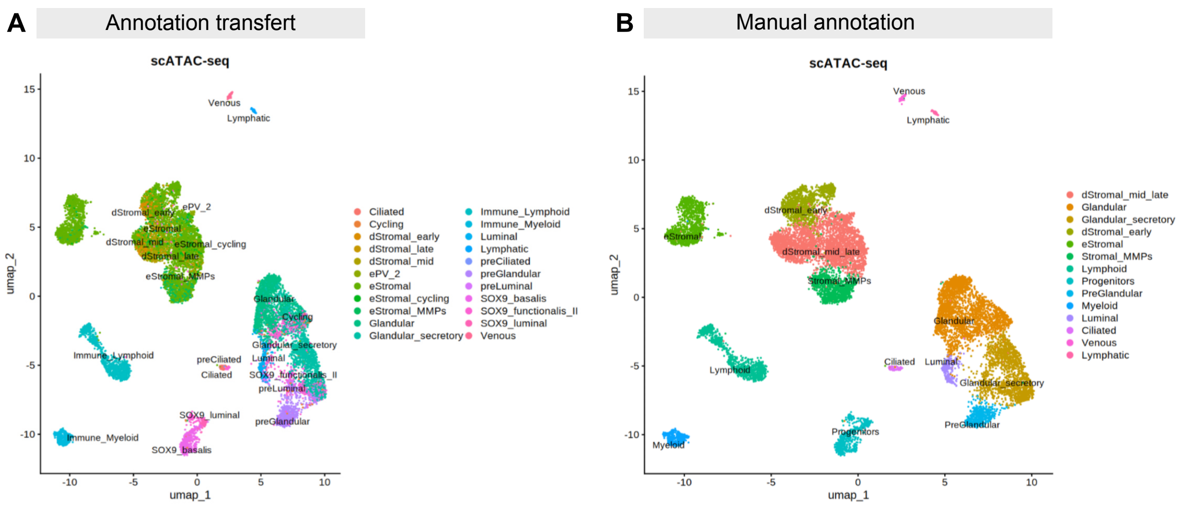

**Figure S6: Annotation of endometrial snATAC-seq from (Vrljicak et al. 2023) by transfer of snRNA-seq annotation from the HECA atlas labels.**

(A) UMAP of snATAC-seq cell type prediction using label transfer from the HECA atlas. (B) Annotation of snATAC-seq dataset used to call peaks.

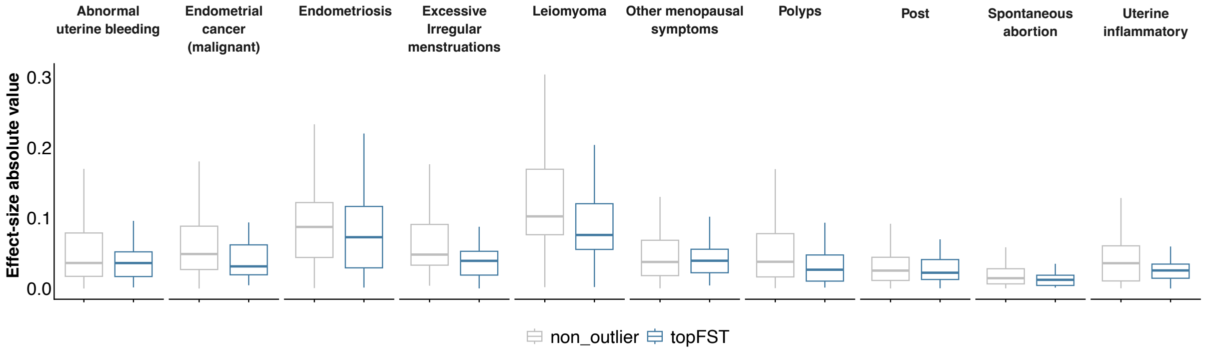

**Figure S7: Effect size differences between uSNPs with outlier Fst values and non-outlier uSNPs.**

Effect sizes correspond to the absolute value of the effect sizes of uSNPs for each uterine disorder. A one-way ANOVA test was performed to assess differences in group means. No significant difference was detected (two-way ANOVA, p-value = 0.44), indicating that the effect sizes between the two groups do not vary significantly.

**Supplementary Tables** (Excel file unless noted)

**Table S1:** List of GWAS phenotypes included from FinnGen

**Table S2:** Genetic correlations estimated by LDSC

**Table S3:** Summary statistics of JASS multi-trait association analysis using FinnGen GWAS, *available on Zenodo: https://doi.org/10.5281/zenodo.19183384*

**Table S4:** List of GWAS phenotypes included from PanUKB

**Table S5:** Replication analysis by directionality of effect sizes

**Table S6:** Summary statistics of JASS multi-trait association analysis using PanUKB GWAS, *available on Zenodo: https://doi.org/10.5281/zenodo.19183384*

**Table S7:** Regulatory annotation of uSNPs

**Table S8:** Gene ontology analysis

**Table S9:** Effect sizes across uSNPs lead variants grouped into clusters

**Table S10:** Prioritized uSNPs with evolutionary metrics annotations
